# Iron associated lipid peroxidation in Alzheimer’s disease is increased in lipid rafts with decreased ferroptosis suppressors, tested by chelation in mice

**DOI:** 10.1101/2023.03.28.534324

**Authors:** Max A. Thorwald, Jose A. Godoy-Lugo, Gilberto Garcia, Justine Silva, Minhoo Kim, Amy Christensen, Wendy J. Mack, Elizabeth Head, Peggy A. O’Day, Bérénice A. Benayoun, Todd E. Morgan, Christian J. Pike, Ryo Higuchi-Sanabria, Henry Jay Forman, Caleb E. Finch

## Abstract

Iron-mediated cell death (ferroptosis) is a proposed mechanism of Alzheimer’s disease (AD) pathology. While iron is essential for basic biological functions, its reactivity generates oxidants which contribute to cell damage and death. To further resolve mechanisms of iron-mediated toxicity in AD, we analyzed postmortem human brain and ApoEFAD mice. AD brains had decreased antioxidant enzymes, including those mediated by glutathione (GSH). Subcellular analyses of AD brains showed greater oxidative damage and lower antioxidant enzymes in lipid rafts, the site of amyloid processing, than in the non-raft membrane fraction. ApoE4 carriers had lower lipid raft yield with greater membrane oxidation. The hypothesized role of iron to AD pathology was tested in ApoEFAD mice by iron chelation with deferoxamine, which decreased fibrillar amyloid and lipid peroxidation, together with increased GSH-mediated antioxidants. These novel molecular pathways in iron mediated damage during AD.

**Graphical Abstract:** 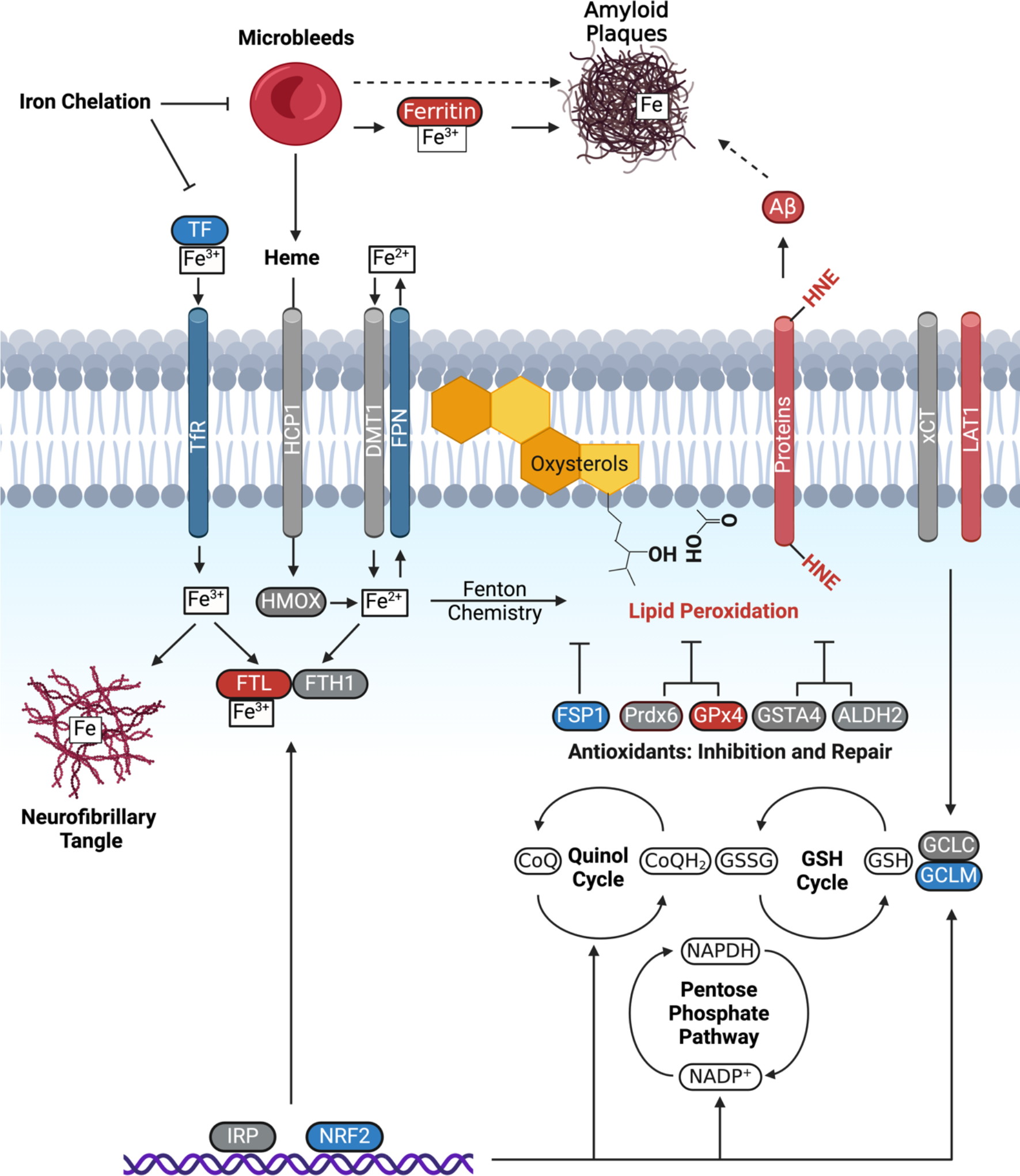

Hypothesis: AD brain lipid peroxidation is driven by increased brain iron and decreased antioxidant defenses. Schema shows proteins that mediate iron metabolism in relation to lipid peroxidation (HNE) and antioxidant defenses in prefrontal cortex. AD-associated increase (red), decrease (blue), or no change (grey), relative to cognitively normal elderly controls. Aβ; amyloid beta, ALDH2; alcohol dehydrogenase, APP; amyloid precursor protein, DMT1; divalent metal transporter 1; FPN, ferroportin; FSP1, ferroptosis suppressor protein 1, which requires the quinol cycle to attenuate lipid peroxidation; FTH1, ferritin heavy chain; FTL; ferritin light chain; GCLC, glutathione cysteine ligase catalytic subunit; GCLM, glutathione cysteine ligase modulator; GPx4, glutathione peroxidase 4; GSH, glutathione; GSSG, glutathione disulfide; GSTA4, glutathione S-transferase A4; HMOX; heme oxygenase; IRP, iron regulatory protein; LAT1, large neutral amino acid transporter 1; LOOH, Lipid hydroperoxides; Nrf2, Nuclear factor erythroid 2-related factor 2; Prdx6, peroxiredoxin 6; TF, transferrin, TfR; Transferrin receptor; xCT, cysteine-glutamate antiporter.

## INTRODUCTION

Postmortem AD neuropathology is traditionally measured using two protein-aggregation related biomarkers: accumulation of amyloid plaques and neurofibrillary tangles (NFT). However, less than 35% of terminal cognitive decline was attributable to these neuropathological markers and was weakly correlated with cognitive status in two large longitudinal studies^1,2^. We suggest new mechanisms for the role of iron in AD neuropathology mediated by glutathione (GSH) and iron export.

Ferroptosis was first designated for cell death *in vitro* from inhibition of cystine glutamate importer (xCT) essential for GSH biosynthesis by quinazoline erastin^3^. GSH is used body wide as an essential substrate for the enzymatic reduction or removal of peroxides, HNE, and xenobiotics. The combination of iron-mediated oxidative damage and its impaired clearance due to GSH deficiency are considered in ferroptosis. Most recently, glutathione peroxidase 4 (GPx4) was identified as a key inhibitor of ferroptosis^4^ due to its role in reducing phospholipid hydroperoxides^5^, which distinguishes it from the other seven GPx family members. GPx4 uniquely interacts with oxidized phospholipids and sterols *within* cell membranes for enzymatic reduction by GSH^6^. We also analyze the GSH-dependent membrane repair protein peroxiredoxin 6 (Prdx6), from a different enzyme family implicated in ferroptois^7^. Additionally we consider the glutathione cysteine ligase modifier (GCLM) subunit critical for GSH homeostasis^8^,. While these studies have expanded the number of proteins in iron-mediated oxidative and repair, other ferroptotic proteins merit consideration in future studies^8,9^.

Iron was first linked to AD in 1953 by Louis Goodman’s evidence for iron within senile plaques and neurofibrils by Prussian blue histochemistry^10^. In the last two decades, associations of iron with AD were strengthened by documentation of increased cerebral microbleeds and decreased blood brain barrier (BBB) integrity, which increase erythrocytes in brain parenchyma. Microglial digestion of intravasated blood yields hemosiderin deposits, consisting of aggregated iron and iron-containing proteins^11^. The accumulation of redox active iron can result in oxidative damage with lipid peroxidation, which are elevated in AD cerebrospinal fluid^12^ and brain^13^. Excessive iron-mediated oxidative damage is implicated in neurodegeneration and cell death. These complex iron-mediated processes are commonly termed ferroptosis, and are extensively studied in cancer and other diseases of aging^14^.

While ferroptosis is implicated in AD pathology^15–17^, there is limited data on human brain for ferroptotic proteins^18–20^ or tissue iron^18,21^. The largest study to date on AD brains showed HNE was increased and varied changes in five proteins that mediate iron storage or transport^18^. This and other reports are limited by the small sample size of 7 or fewer brains each, without identifying sex or ApoE allele. We measured these and twenty additional proteins related to iron metabolism, antioxidants, and amyloid pathology. A potential confound of prior studies is residual blood from unperfused brains. Blood contains several fold higher levels of heme-iron, iron bound proteins, and GSH-utilizing enzymes than brain. Experimental knockdown of GPx4 in mice caused neuron loss supporting the concept of ferroptosis^22^, which is cited in reviews as support for ferroptosis in AD. While GPx4 has an essential role in neuronal survival, AD brains have not been examined for GPx4 or other antioxidants processes relevant for ferroptosis. Moreover, xenografting human AD neurons into mice caused necrotic cell death, but not apoptosis or ferroptosis^23^, while a murine model of acute kidney disease intertwinement of necroptosis and ferroptosis^24^. The mechanisms linking specific cell death processes during AD needs further clarification.

To further resolve the role of iron in AD, we comprehensively examined human postmortem tissue for proteins implicated in initiation or inhibition of ferroptosis. A large set of brains from cognitively normal and AD was examined with equal sex and known Apolipoprotein E (ApoE) alleles. The prefrontal cortex was compare with the ‘AD resistant’ cerebellum^25^ for multiple proteins of iron metabolism and antioxidant defense. The lipid raft microdomain, where the bulk of amyloid precursor protein (APP) is processed for β-amyloid (Aβ) peptides, was analyzed for pathways relevant to ferroptosis. We tested iron dependent mechanisms in a mouse model of AD by chelation with deferoxamine (DFO) used in clinical trials for AD^26^. These findings document the loss of antioxidant enzymes in lipid rafts and associated increases in lipid peroxidation.

## RESULTS

### Minimization of residual blood in postmortem brain

Studies of iron in postmortem tissues contain intravascular blood, rich in iron proteins. In rodent studies, blood can be removed by cardiac perfusion with heparinized saline, which we show decreased heme iron in the cerebral cortex by 35% (**Table 1A**). Because perfusion is not feasible for thawed human cerebral cortex, we developed a washing procedure to minimize adherent and entrapped vascular blood. Thawed brain was minced into 2 mm cubes, followed by gentle washing with isotonic buffer (PBS). ‘Washed’ tissues had 65% less heme than the non-perfused, displaying less heme even when compared to perfused samples which were not washed (**Table 1A)** suggesting most heme is from residual blood on tissue rather than in the vasculature.

**Table 1.** Heme and Iron content of prefrontal cortex. Heme and total iron in prefrontal cortex were assayed for effects of washing to minimize residual intravascular blood. **A**) mouse (C57BL/6J, 6 mo; n=5-6); **B**) human (AD; n=24) and controls (65+ years; n=24) prefrontal cortex (Brodmann regions 8-10); **C)** Iron^21,28,29^ and ferritin^29–31^ in brain, blood, serum, and CSF. The ferritin complex in erythrocytes and tissue include FTL and FTH1. Isotonic phosphate buffered saline (PBS; pH 7.4) was used for aortic perfusion of mouse and for washing of minced mouse and human brain. 30mg prefrontal cortex was minced into 2 mm cubes and ‘washed’ by gentle vortexing in PBS. Heme and α- hemoglobin per mg tissue were assayed spectrophometrically relative to unwashed brain (**Fig. S3**). Total iron levels from Ashraf^33^ were estimated as Fe/g protein, assuming 1 g protein/12 g wet weight brain. Brain iron and ferritin by compartment for human are >95% below blood levels, assuming 1 ml volume is equivalent to 1-gram wet brain. Significance by one way ANOVA (mouse) with Tukey’s posthoc, or 2-tailed t-test (human): *p<0.05, **p<0.01, ***p<0.001, ****p<0.0001.

|  | Non-perfused | Perfused PBS | Perfused PBS +<br>Heparin | Non-Perfused +<br>Washed |
| --- | --- | --- | --- | --- |
| Heme/mg wet brain | 3.1 ± 0.2 | 2.1 ± 0.2* (-32%) | 2.0 ± 2.3** (-35%) | 1.3 ± 0.2*** (-68%) |

**B. Human**
| Source | Unwashed | Washed |
| --- | --- | --- |
| Total iron, ug/g wet brain | 45 ± 4.9 | 26 ± 1.4 ****, -45% |
| Ayton et al. 2021<sup>21</sup> | ~55 | N/A |
| Ashraf et al. 2020<sup>18</sup> | ~290 | N/A |
| Heme/mg | 7.1 ± 0.4 | 3.6 ± 0.2, ****, -50% |
| a-Hemoglobin, ug/mg | 1,644 ± 131 | 815 ± 61, ****, -50% |
| Ferritin light chain (RFUs) | 235 ± 17 | 100 ± 9, **** -60% |

**Table 1:** Heme and total iron in prefrontal cortex were assayed for effects of washing to minimize residual intravascular blood. **A)** mouse (C57BL/6J, 6 mo; n=5-6); **B)** human (AD;
| Compartment | Total iron | Ferritin |
| --- | --- | --- |
| Brain, unwashed | 50 ug/g wet brain | 0.135 ug/g |
| Blood | 3,500 ug/mL | 3.8-14 ug/mL |
| Serum | 1.3 ± 0.14 ug/mL | 0.011-3.4 ug/mL |
| CSF | 0.061 ± 0.006 ug/mL | 0.012 ug/mL |

For human cortex (**Table 1B)** washing decreased levels of heme by 50%. As expected, washing reduced total iron levels significantly in the cortex (**Table 1B, Fig. S1A**). Importantly, total iron levels in unwashed samples were similar to those in previous studies with an increase observed with AD, suggesting that iron in previous reports^18,21,27^ were possibly influenced by residual blood.

Total aluminum, copper, and zinc, which generally have low levels in the blood, were not altered by washing (**Fig. S1B-D**), supporting that our washing protocol was specific to removing blood.

We further validated our washing protocol by centrifugation of the residual PBS was from tissues. The remaining supernate contained heme levels equivalent to the difference between washed and unwashed tissue (**Fig. S2A**). Serial washes were performed which showed diminishing decreases in heme concentrations after the second wash (**Fig. S2B**), for these reasons tissues were washed twice for the remaining measurements. a-hemoglobin and ferritin light chain, other markers for erythrocytes, all of which showed decreased levels greater than 50% in our washed samples (**Fig. S2C-F**). While our data suggest that some residual blood still exists in samples after washing, our protocols eliminated blood contaminants by half. Therefore, for the entirety of this study, all analyses were performed on washed samples to eliminate this confounding variable.

### Lipid peroxidation increased in AD

First, we measured levels of lipid peroxidation, the oxidation of lipids by free radicals (**Fig. 1A**), previously shown in other AD tissues^34^ which are a main component of ferroptosis. Oxidative damage was assessed in the prefrontal cortex with two protein adducts, HNE and 3-nitrotyrosine (NT). AD samples exhibited a significant increase in both HNE and NT compared to controls (**Fig. 1B-C**), suggesting increased lipid peroxidation is correlated with AD pathology. To determine whether these changes were associated with ApoE status or sex, we next measured HNE and NT in ApoE3 vs. ApoE4 carriers and found that lipid peroxidation was higher in ApoE4 carriers, consistent with previous reports that ApoE4 carriers are at higher risk for pathology (**Fig. 1D**). When separated by sex, females displayed an increase in HNE, but males only showed a trend, but not significant, increase in HNE, also consistent with previous reports that females display worse AD pathology than males^35^ (**Fig. 1E**).

**Figure 1:**
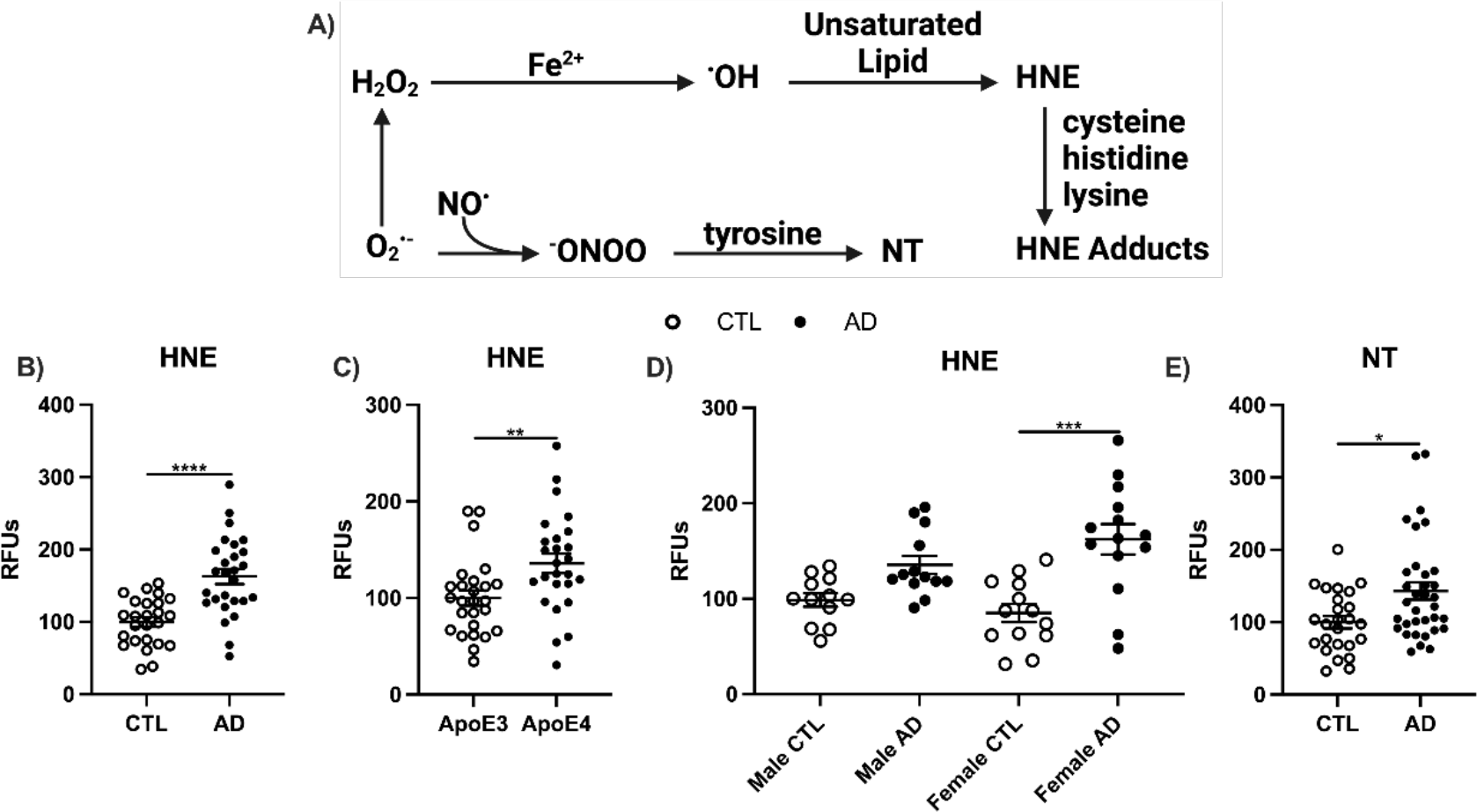
Oxidative damage (HNE, NT) to prefrontal cortex from age matched AD (n=27) and cognitively normal controls (CTL; n=25). **A)** Schematic representing oxidative damage to lipids and proteins. Hydroxyl radicals (.OH) are produced during oxidation of Fe^2+^ or Cu^2+^ (‘Fenton chemistry’) and may oxidize adjacent proteins, lipids, or nucleic acids. HNE can alter proteins via Michael addition products through its double-bonded carbons, or by forming Schiff bases through its aldehyde moiety. The bottom line shows the iron-independent oxidative pathway to form peroxynitrite ^-^ONOO from O2^•–^ and .NO, which can then oxidize tyrosine to NT. Aggregated human Aβ42 stimulated formation of peroxynitrite which is neurotoxic in vitro^36^. Non-heme iron also mediates lipoxygenase activity^37^. Dot blots as relative fluorescent units (RFUs) for HNE presented as **B)** cognitively normal vs AD, **C)** ApoE allele, or **D)** sex and **E)** NT. Significance, 2-tailed t-test (B-C, E), one-way ANOVA with Tukey’s posthoc test (D): *p<0.05, **p<0.01, ***p<0.001, ****p<0.0001.

Next, we examined changes in lipid peroxidation by brain region. First, we confirmed that like cortex samples, washed cerebellum only displayed a mild increase in heme when comparing AD to control samples (**Extended Data** Fig. 1A). Cerebellum samples did display increased HNE with AD compared to control, although to a lesser extent than in the cortex (**Extended Data** Fig. 1B). In addition, no difference was observed in NT when comparing AD to control samples in the cerebellum, although cortex samples do show a difference in NT between AD and control (**Extended Data** Fig. 1C). The most striking was the loss of association of ApoE status on lipid peroxidation in the cerebellum (**Extended Data Fig. 1**D), and the reversal of association of sex: males displayed higher levels of lipid peroxidation in AD samples, whereas no difference was seen in females in the cerebellum (**Extended Data Fig. 1**E). Finally, to confirm that changes in lipid peroxidation correlate with “canonical” markers for AD pathology, we confirmed that our AD cortex and cerebellum samples display an increase in Aβ fibrils (**Extended Data Fig. 1**F) Altogether, these data suggest that the increased lipid peroxidation is associated with fibrillar amyloid. In addition, although the cerebellum does have milder AD pathology compared to the cortex in terms of lipid peroxidation, they are not completely devoid of pathology. Importantly, the association of ApoE status and sex on lipid peroxidation seems to be brain-region specific.

### Iron protein metabolism is decreased in AD

While washed brain tissues did not have increases in total iron, many studies have shown that there is increased infiltration of blood cells into the brain during progression of AD due to microbleeds and loss of BBB integrity. Thus, although for biochemical assays, it is important to remove blood to differentiate between contribution of iron and iron-related factors in the blood vs. brain tissue, it is important to remember that in the context of an intact human brain, blood-contribution of iron is still an important driver in pathology. Therefore, we next sought to determine the changes in iron metabolism in AD brains, which could be driven by blood cell infiltration. Brain cellular iron is imported by the divalent metal transporter 1 (DMT1) together with transferrin (TF) and its receptor TfR. The levels of TF and TfR differ by cell type^19^. Heme iron is imported by heme carrier protein 1 (HCP1) and degraded by hemeoxygenases (HMOX1 or HMOX 2). Intracellular ferric iron is stored in complex with ferritin light chain (FTL) after oxidation of ferrous iron (Fe^2+^) by ferritin heavy chain (FTH1). Export of ferrous iron is mediated by ferroportin (FPN).

As expected, we saw that numerous proteins involved in iron were depleted in AD brains. First, iron-transport proteins TF and its receptor TfR, were decreased in AD brains compared to control (**Fig. 2A-B**). Although we did not observe a statistically significant decrease in DMT1, there is a trend for a decrease in AD brains (**Fig. 2C**), and FPN also showed significant reduction (**Fig. 2D**).

**Figure 2:**
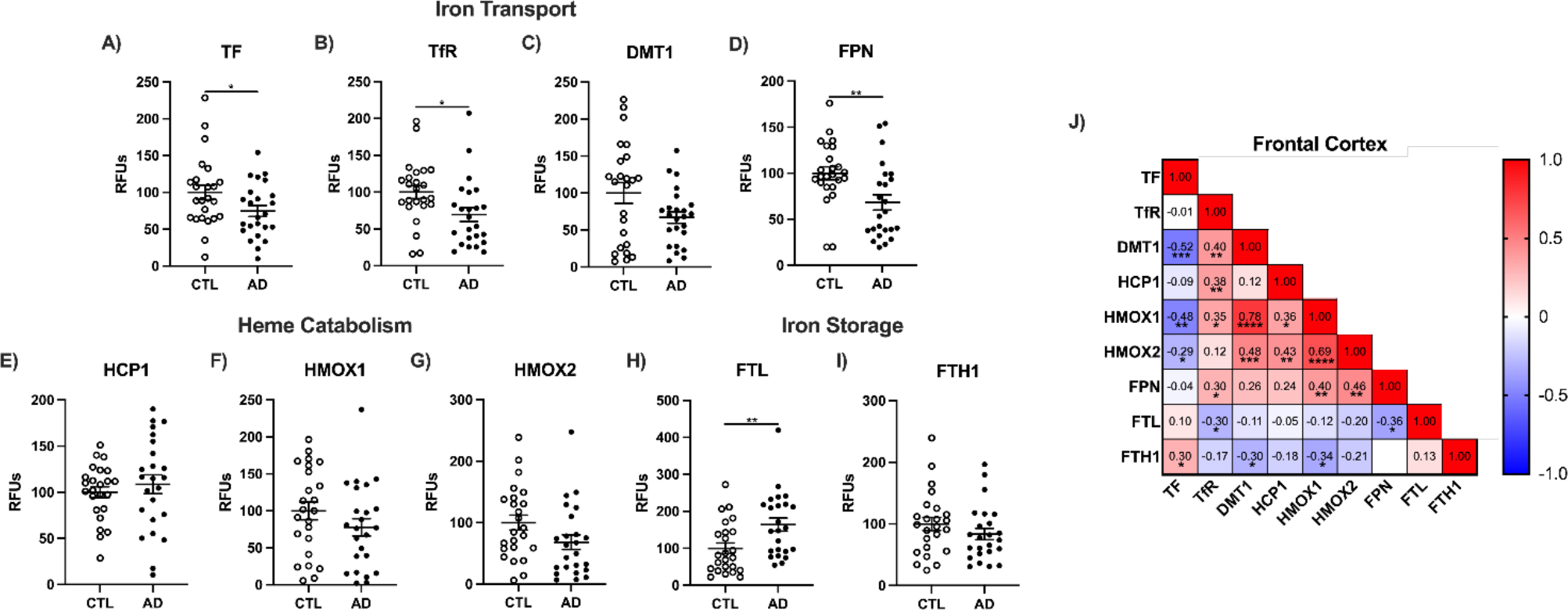
Iron transport and metabolism in AD prefrontal cortex (n=24) compared to cognitively normal age-matched controls (n=24). Relative fluorescent units (RFUs) from Western blots. **A)** transferrin (TF), **B)** transferrin receptor (TfR/CD71), **C**) divalent metal transporter 1 (DMT1), **D)** ferroportin (FPN), **E)** heme carrier protein 1 (HCP1), **F)** hemeoxygenase 1 (HMOX1), **G)** heme oxygenase 2 (HMOX2), **H)** ferritin light chain (FTL), **I)** ferritin heavy chain 1 (FTH1), **J)** Correlation matrix for iron metabolism in clinical AD. Significance, 2-tailed t-test (A-I), Spearman correlation (J): *p<0.05, **p<0.01, ***p<0.001, ****p<0.0001.

Interestingly, we did not observe major differences in heme catabolism (**Fig. 2E-G**). In addition, there was a significant increase in the iron storage protein, FTL (**Fig. 2H**), although FTH1 did not show any difference between AD and control samples (**Fig. 2I**). Relationships for iron import/export include negative correlations between iron storage protein FTL with iron import receptor TfR (r=-0.30, p=0.04) and export protein by FPN (r=-0.36, p=0.01; **Fig. 2J**). ApoE4 carriers had 2-fold higher FTL than ApoE3 in prefrontal cortex (**TableS3**). Interestingly, AD cerebellum had dramatically different changes in iron metabolism proteins compared to the cortex. The only iron transport protein that increased in AD cerebellum compared to control samples was TF (**Extended Data Fig. 2**A), with all eithers either displaying no change or a significant decrease (**Extended Data Fig. 2**B-D). Similar to most iron transport proteins, proteins involved in heme catabolism were also largely unchanged, with only HMOX1 showing a slight decrease (**Extended Data Fig. 2**E-G). Finally, iron storage proteins also did not display any difference in AD cerebellum (**Extended Data Fig. 2**H-I).

Overall, our data show that even though total iron levels do not change in the brain, there are significant differences in proteins related to iron metabolism. These changes seem to be primarily limited to the cortex, which suggests that the cortex is more susceptible to changes to iron exposure when compared to the cerebellum.

### GSH synthesis is decreased with AD

Lipid peroxidation is driven by the loss of redox homeostasis. The tripeptide GSH conjugates to HNE for removal by several enzymes. GSH is also essential to reduce peroxides by the GPx enzyme and Prdx families. GPx4 and Prdx6 are the only known lipid hydroperoxidases capable of reducing oxidized phospholipids. These enzymes are critical in safeguarding against lipid peroxidation and ferroptosis. All other GPx family members also detoxify hydroperoxides using GSH, with GPx1 being the most abundant GPx family member in brain^38^. Shockingly, we saw an increase in GPx4 levels in AD prefrontal cortex when compared with controls (**Fig. 3A**). However, total GPx4 activity was not changed (**Fig. 3B**), suggesting that although enzyme levels may be higher, the functional output of these hydroperoxidases are unchanged. In addition, GPx1 levels and activity, as well as levels of Prdx6 were unchanged in the AD cortex (**Fig. 3C-E**). Finally, ferroptosis suppressor protein 1 (FSP1), which also protects against lipid peroxidation through the quinol cycle in a GSH- independent manner did show a significant reduction in the AD cortex (**Fig. 3F**).

**Figure 3:**
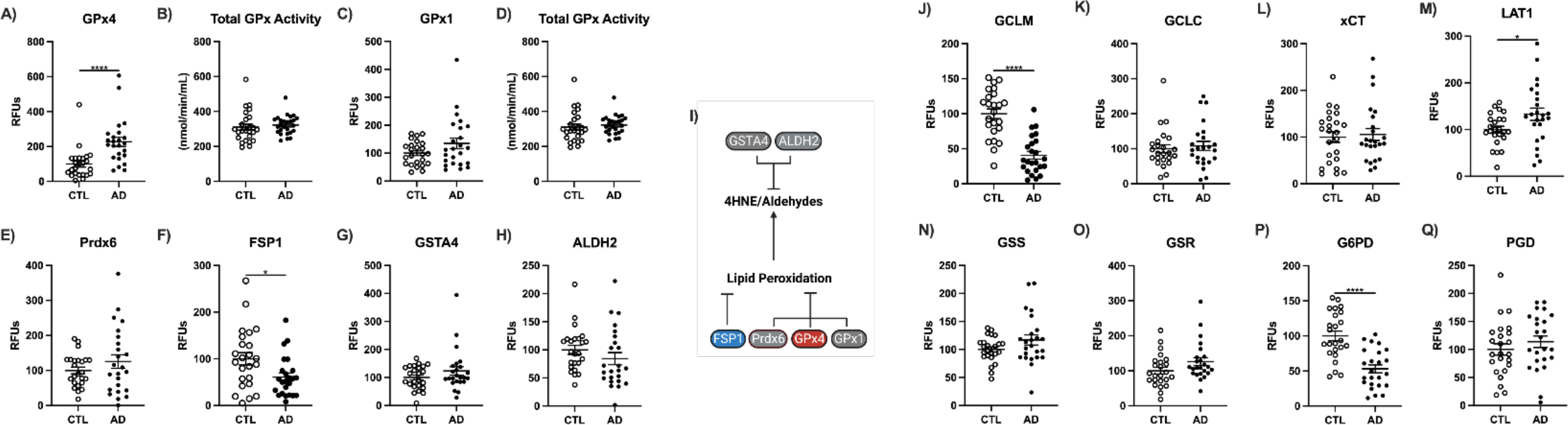
Enzymes that repair or mitigate lipid peroxidation and facilitate GSH production. Human prefrontal cortex Western blots and enzymatic activity in whole tissue lysate for **A)** GPx4, **B)** GPx4 activity, **C)** GPx1, **D)** total GPx activity, **E)** Prdx6, **F)** FSP1, **G)** GSTA4, and **H)** ALDH2. **I)** Schema of enzymatic repair of lipid peroxidation. GSH cycle enzyme levels by Western blots: **J)** GCLM, **K)** GCLC, **L)** xCT/SLC7A11, **M)** LAT1, **N)** GSS, **O)** GSR, **P)** G6PD, and **Q)** PGD. Significance, 2-tailed t-test (A-H, J-Q): *p<0.05, ****p<0.0001.

To address the discrepancy between increased lipid peroxidation observed in AD brains and the lack of changes seen in detoxification enzymes, we next sought to determine whether there were defects in enzymes involved in GSH synthesis (**Fig. 3I**). The glutathione S-transferases (GST) conjugates GSH with diverse metabolites and xenobiotics. Biosynthesis of GSH depends on cystine import via the proteins xCT (SLC7A11) and LAT1 (SLC3A2/CD98). Imported cystine is rapidly reduced to cysteine by the intracellular reducing environment^3^. The rate limiting step in GSH synthesis^39^ is formation of γ-glutamylcysteine, catalyzed by glutamate cysteine ligase, with two subunits GCLC (catalytic) and GCLM (modulatory); the binding of GCLM to GCLC accelerates GSH synthesis four-fold. Next, GSH forms from addition of glycine to γ-glutamylcysteine by glutathione synthetase (GSS). Since GPx4 and Prdx6 activities are dependent on GSH, a decreased total pool of GSH would also increase lipid peroxidation, even if GPx4 and Prdx6 protein levels remain unchanged. GSH itself could not be measured due to its rapid oxidation upon death and associated artifacts, which is impossible to prevent in human postmortem tissues^40^. In the prefrontal cortex, the glutamate-cysteine ligase subunits were differentially altered by AD: the GCLM modulatory subunit was 60% lower, while GCLC the catalytic subunit was unaltered (**Fig. 3J,K**). LAT1 (L-type amino acid importer transporter) increased 33% in AD, while xCT did not differ, contrary to the 20% increase reported by Ashraf *et al.* 2020^18^ (**Fig. 3L,M**). Two enzymes of the GSH cycle were not altered in AD cortex: GSS, the final step in GSH synthesis (**Fig. 3N**) and glutathione reductase (GSR) (**Fig. 3O**), which utilizes NADPH to reduce glutathione disulfide back to GSH.

Related metabolic cycles also showed selective AD changes. For the pentose phosphate pathway, G6PD was decreased by 45% in AD relative to controls, while phosphogluconate dehydrogenase (PGD) was unchanged (**Fig. 3P,Q**). These enzymes oxidize glucose to generate NAPDH required by GSR for oxidized GSH reduction (GSSG). Two groups of enzymes had correlated levels: G6PD and PGD for NADPH-production were positively correlated with GSR (both r=0.31, p=0.03), as were the amino acid importer LAT1 and cystine importer xCT (r=0.49, p<0.001; not shown). Collectively, the decreases in GCLM and G6PD in the AD prefrontal cortex suggest reduced GSH levels, which contributes to the increased lipid peroxidation, despite a lack of change in GPx4 or antioxidant enzymes.

Unlike the cortex, AD cerebellum showed more changes to lipid hydroperoxidases. GPx4 levels decreased in AD cerebellum (**Extended Data Fig. 3**A), although total activity of GPx4 was unchanged (**Extended Data Fig. 3**B). In addition, although GPx1 levels and activity were unchanged like AD cortex (**Extended Data Fig. 3**C-D), Prdx6 levels were lower in AD cerebellum (**Extended Data Fig. 3**E). Finally, consistent with changes in AD cortex, FSP1 levels were also decreased in AD cerebellum (**Extended Data Fig. 3**F). Overall, AD cerebellum displayed more changes antioxidant enzymes compared to the cortex, in addition to showing a similar decrease in the GSH synthesis enzyme, GCLM (**Extended Data Fig. 3**J). Although other GSH synthesis enzymes were largely unchanged, apart from GSS which increased in the cerebellum (**Extended Data Fig. 3**K-Q).

### Lipid peroxidation in lipid rafts is increased and antioxidant defense is diminished in AD

Since measurements of whole-cell protein levels of antioxidant defense enzymes were inconclusive, we next examined the local environment of subcellular lipid rafts (**Fig. 4A**), the hub of signal transduction and where APP is processed. Lipid raft isolations were verified and compared with traditional ultracentrifugation methods^41^. First, we characterized which of the eight GPx isoforms are present in lipid rafts and detected only GPx4 and GPx1 (**Fig. S3**). Prdx6 was not detected. AD lipid rafts had significantly lower levels of the membrane protective enzymes GPx4 and GPx1, as well as decreased activity of both GPx4 and GPx1 (**Fig. 4B-E**). In addition, we observed lower levels of ALDH2 which clears free HNE and the Aβ clearance proteins, ApoE and LRP1 in AD lipid rafts (**Fig. 4F-H**). Importantly, the decreases in antioxidant protection and activity in the AD cortex were further substantiated with increased lipid raft oxidation using HNE and NT (**Fig. 4I-J**), paralleling increases in total tissue (**Fig 1B-E**). This increase in lipid peroxidation is associated with the ApoE4 allele, which exhibited higher levels of HNE in lipid rafts (**Fig. 4K**) but had no associations with sex as both males and females showed similar increases (**Fig. 4L**). Overall, these data suggest that measurements of oxidative stress, particularly linking changes to antioxidant defense and lipid peroxidation, are most robust in the lipid rafts for AD samples. Collectively, our data strongly correlate the loss of antioxidant enzymes in rafts with lipid oxidative damage in AD cortex.

**Figure 4:**
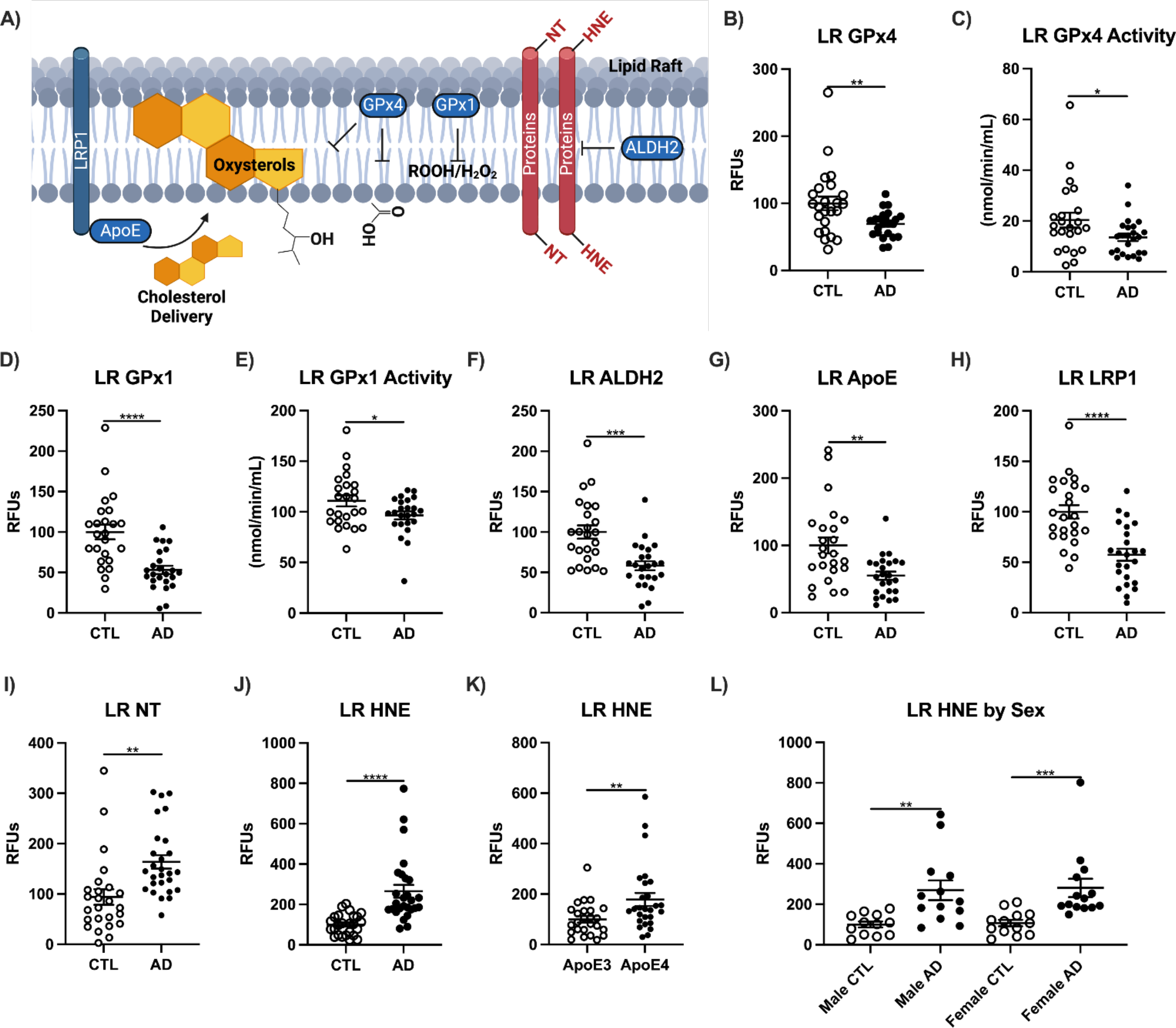
The lipid raft membrane fraction has increased damage and reduced antioxidant defense during AD. **A)** Schema of lipid raft damage, protective mechanisms, and cholesterol shuttling. Western blot data and enzymatic activity **B)** GPx4, **C)** GPx4 activity, **D)** GPx1, **E)** GPx1 activity, **F)** ALDH2, **G)** ApoE, **H)** LRP1. Dot blots for **I)** NT, and HNE presented as **J)** CTL vs AD, **K)** ApoE allele, and **L)** sex. Significance, by 2-tailed t-test (B-K) or one-way ANOVA with Tukey’s posthoc (L): *p<0.01, **p<0.01, ***p<0.001, ****p<0.0001.

Next, we performed similar lipid raft measurements in the cerebellum and found that like the AD cortex, AD cerebellum exhibited a significant decrease in GPx4 and GPx1 levels. However, unlike the cortex, GPx4 and GPx1 activity were not decreased in AD cerebellum (**Extended Data Fig. 4**A**- E**). Although ALDH2 levels were unchanged, ApoE and LRP1 were significantly decreased in AD cerebellum (**Extended Data Fig. 4**F-H). Similar to what was observed in whole-cell lipid peroxidation, AD cerebellum lipid rafts displayed no change in NT, but higher levels of HNE, which is consistent with our previous conclusion that in regard to lipid peroxidation, cerebellum samples display lower AD pathology than the cortex. Similar to the cortex, the cerebellum also showed higher lipid peroxidation in ApoE4 carriers (**Extended Data Fig. 4**L) with no observable difference between males and females (**Extended Data Fig. 4**M).

The non-raft membrane fraction from these same samples was also examined, which unexpectedly showed increased oxidative damage for NT but not HNE (**Extended Data Fig. 4**N-Q). No ApoE allele differences were observed for the non-raft membrane. Non-raft membrane GPx4 did not differ by AD (**Extended Data Fig. 4**R). GPx4 in non-membrane fraction was 50% lower below LR from the same tissues (**Extended Data Fig. 4**S), suggesting more localization of GPx4 to lipid rafts. These data suggest some membrane domains are selectively vulnerable to oxidation.

### Experimental iron chelation improves pathology in AD mouse models

Thus far, our data provides strong correlative evidence that dysfunction of iron homeostasis can drive lipid peroxidation, which is exacerbated in lipid rafts due to a loss of antioxidant enzymes in AD brains. However, one major limitation of studies with postmortem brain tissue is the inability to experimentally test causality. Therefore, we performed chelation by DFO to evaluate how removal of iron can impact AD pathology in the EFAD mouse model. DFO was chosen over other chelators because of its use in AD patients^26^ and experimental studies with [H^3^]DFO show it crossed the BBB, yielding 3-fold higher levels per gram in the brain than blood and kidney^42^. In our experimental paradigm, female EFAD mice received DFO in two modes: by lab chow for two weeks or by intraperitoneal injection (IP) twice daily for one week (see extended data for comparison of DFO modes).

First, we confirmed that DFO treatment ameliorated canonical markers of AD: specifically, we found that DFO-treated mice displayed a significant reduction in fibrillar Aβ (**Fig. 5A, Extended Data** Fig. 5A), and a significant increase in soluble Aβ peptides (**Fig. 5B-C, Extended Data** Fig. 5B-C).

**Figure 5:**
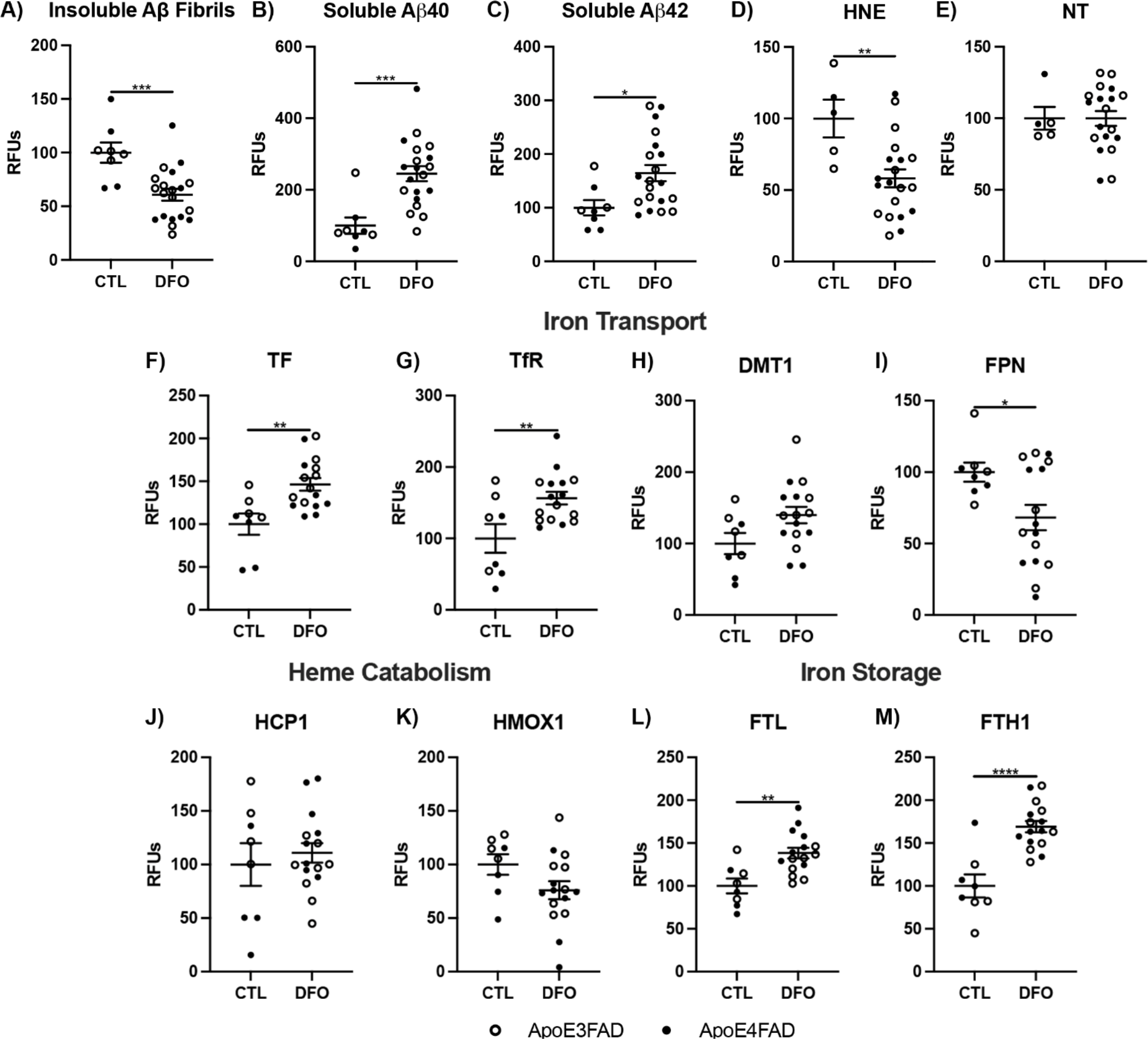
DFO chelation of EFAD mouse cortex: **A)** insoluble Aβ fibrils, **B)** soluble Aβ40, **C)** soluble Aβ42, **D)** HNE, **E)** NT, **F)** TF, **G)** TfR, **H)** DMT1, **I)** FPN, **J)** HCP1, **K)** HMOX1, **L)** FTL, and **M)** FTH1. Significance, 2-tailed t-test: *p<0.05, **p<0.01, ***p<0.001, ****p<0.0001.

Importantly, we observed a significant reduction in lipid peroxidation using HNE in DFO-treated mice, and no change in NT which confirms the role of iron in lipid peroxidation (**Fig. 5D-E, Extended Data** Fig. 5D-E). To confirm whether these changes were due to changes in iron handling, we next measured changes in proteins related to iron metabolism. Indeed, we found that many iron transport proteins and iron storage proteins were significantly increased, although no significant changes were seen in heme catabolism proteins (**Fig. 5F-M, Extended Data** Fig. 5F-M).

As previously mentioned, since ferroptosis is induced by a combination of iron overload and/or an increase in oxidative damage, we next measured whether iron chelation through DFO was sufficient to improve redox handling and antioxidant activity in the brain. DFO-treatment resulted in a significant increase in GPx4 and GPx1 levels, with a significant increase in GPx1 activity, but not GPx4 activity, although a trend of increase was seen for GPx4 activity in some animals (**Fig. 6A-D, Extended Data** Fig. 6A-D). In addition, GSTA4, GCLC, and xCT showed a significant increase in DFO-treated animals, although several others (Pdrx6, FSP1, ALDH2, GCLM) showed no difference (**Fig. 6-E-K, Extended Data** Fig. 6E-K). Overall, these experimental studies provide direct evidence that removal of iron and improved iron handling can reduce oxidative damage, improve antioxidant defense, and ameliorate AD pathology.

**Figure 6:**
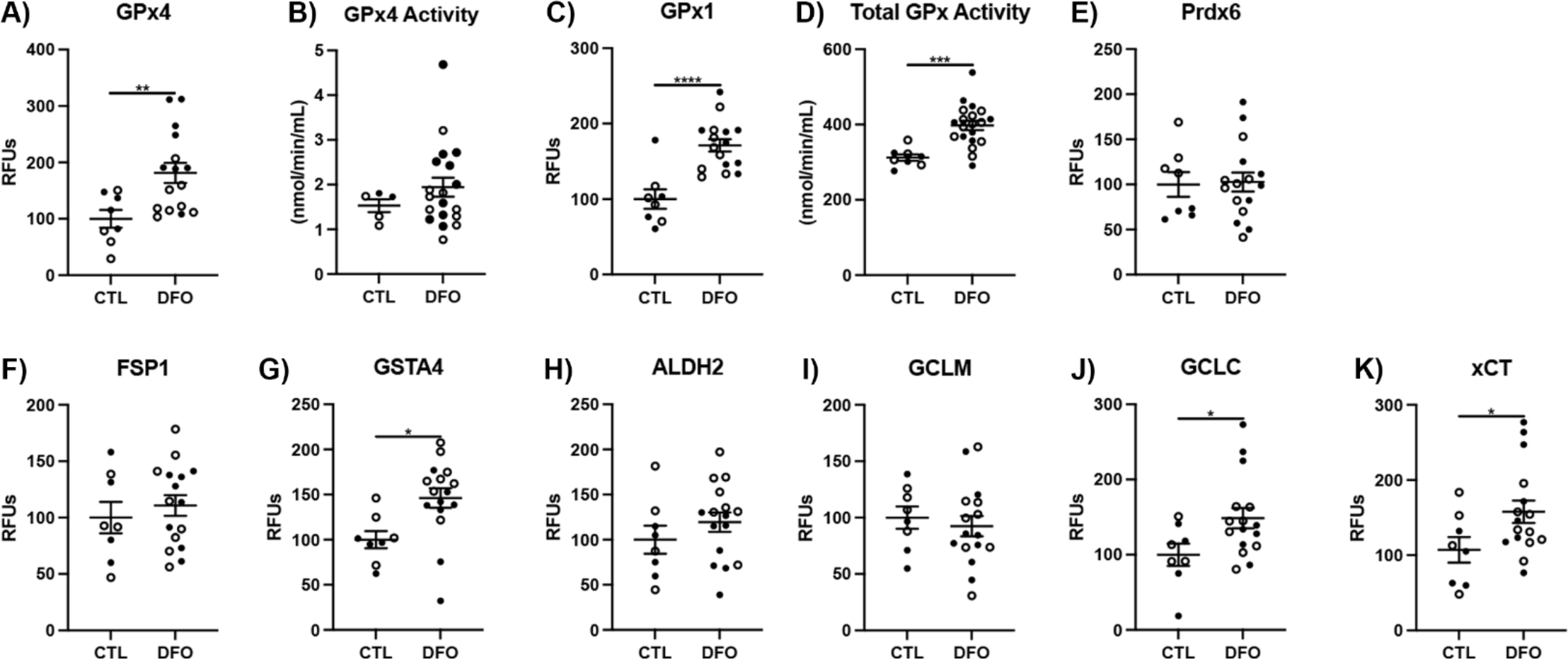
DFO chelation of EFAD mouse cortex. **A)** GPx4, **B)** GPx4 activity, **C)** GPx1, **D)** GPx1 activity, **E)** Prdx6, **F)** FSP1, **G)** GSTA4, **H)** ALDH2, **I)** GCLM, **J)** GCLC, **K)** xCT. Significance, 2-tailed t-test: *p<0.05, **p<0.01, ***p<0.001, ****p<0.0001.

### AD pathogenesis downstream of iron-related oxidative damage is driven through changes in Nrf2 signaling

To determine whether the outcome of iron-related oxidative damage in AD pathology was due to a dysregulation of iron homeostasis directly or through changes in downstream antioxidant pathways, we next measured changes in major transcriptional regulators of iron and oxidative damage handling. Interestingly, we found no major differences in the transcription factors, IRP1 and IRP2, involved in regulation of iron genes in the AD cortex, although a decrease in IRP2 was observed in the AD cerebellum (**Fig. S4A-C**). Importantly, we saw a significant decrease in Nrf2 in the AD cortex, one of the major transcription factors involved in antioxidant defense pathways and iron regulation (**Fig. S4D**). Interestingly, the AD cerebellum did not display a significant decrease in Nrf2. Importantly, the AD cortex also showed higher levels of NF-kB, associated with inflammation known to be elevated during AD (**Fig. S4E**). Like Nrf2, NF-kB did not show any observable differences in the cerebellum. Finally, there was a significant increase in nuclear receptor coactivator 4 (NCOA4) in the AD cortex (**Fig. S4F**) which signals for iron degradation by ferritinophagy^43^. These data were consistent with our other observations where changes in antioxidant defense pathways were more pronounced than changes in iron metabolism in the AD brain. Moreover, consistent with our previous data, the AD cortex displayed more oxidative damage than the cerebellum.

To determine whether this depletion of nuclear Nrf2 correlated with changes in gene expression of genes involved in antioxidant defense, we performed RNA-seq analysis on control-matched and case-matched AD cortex and cerebellum (**Fig. S4G-H**). Although for the purpose of this study, data were pooled into AD vs. control, we ensured that RNA-seq analysis was performed across males, females, and ApoE genotypes, which theoretically can be analyzed across sexes and genotypes in a future study. GO analysis revealed that many differentially expressed genes in the AD samples were those associated with numerous pathways previously implicated in AD pathology, including mitochondrial dysfunction (**Fig. S4I**). Many of these significant GO terms overlapped between the cortex and the cerebellum, although there were also several terms that were not shared, suggesting again that the cerebellum is not fully AD resistant, although the cortex displays signs of higher pathology. As expected, we did not observe major differences in gene expression of iron-related genes. More importantly, we found depletion of several Nrf2 targets in the AD cortex, but not in the cerebellum (**Fig. S4J**), which directly correlates with the decrease in nuclear Nrf2 found specifically in the AD cortex. Transcriptome changes shared only 9 common DEGs between brain regions. CD163, the hemoglobin-haptoglobin scavenger which increases in response to microbleeds was shared between brain regions with AD and was among the top responding mRNAs^44^. Other genes previously associated with AD or dementia were differentially expressed in both brain regions: CHI3L2, FCGBP, GLI1, STAB1, APOC1, MYO7A, and long noncoding RNA JHDM1D-AS1 (**Fig. S4K**). Taken together, our assay of transcriptional pathways supports our current hypothesis that iron toxicity is directly associated with a loss of antioxidant defense in AD pathology.

Consistent with the human data, we also found that DFO treatment of EFAD mice did not increase IRP or NOCA4 but did result in a significant increase in Nrf2 (**Fig. S4L-O**), again suggesting that most pathogenic pathways downstream of iron are through changes in antioxidant defense pathways.

### Neuronal loss in AD is associated with increased ferritin, lipid peroxidation, and diminished antioxidant defenses

To next determine the physiological consequence of iron-mediated oxidative damage to lipids, we measured neuronal loss in AD samples. While neuronal loss dramatically increases with pathology in AD, the relationship between neuronal loss iron or membrane damage in AD is still lacking. Therefore, we analyzed bulk neuronal content in prefrontal cortex and cerebellum using NeuN (neuronal nuclear antigen), presynaptic SYP (synaptophysin1), and postsynaptic PSD95 (postsynaptic density protein 95). Strikingly, we saw a significant depletion of NeuN and PSD95, and a trend for a decrease in SYP in AD cortex compared to control (**Fig. 7A-C**), although no significant changes were seen for any markers in the cerebellum (**Extended Data Fig. 7**A-C). These data also by Braak Score shows a similar trend whereby higher Braak scores correlated with lower NeuN and PSD95 in the cortex, but not the cerebellum (**Fig. 7D-F, Extended Data Fig. D-F**). In addition, neuronal markers varied inversely with cortex tissue levels of heme by Braak score, while NeuN varied inversely with ferritin light chain (**Fig. 7D**). The relationship for NeuN and heme differed by Braak stage, but not by clinical dementia status. These inverse associations give the first direct associations of brain iron markers with brain region-specific neuron loss during AD. The inverse correlation of NeuN with HNE strengthens the relationship of lipid peroxidation to neuron loss. Many of these relationships were weakened or absent from cerebellum, consistent with its reduced neuronal loss (**Extended Data Fig. 7**D-E). However, positive correlations between NeuN and FPN or NeuN and GCLM protein levels were the only shared relationships by both criteria and in cortex and cerebellum. These relationships highlight the role iron export by FPN or GSH production by GCLM for neuronal survival. Overall, these data provide direct evidence that the physiological consequence of increased iron-related oxidative damage results in neuronal cell death, likely through the induction of ferroptosis.

**Figure 7:**
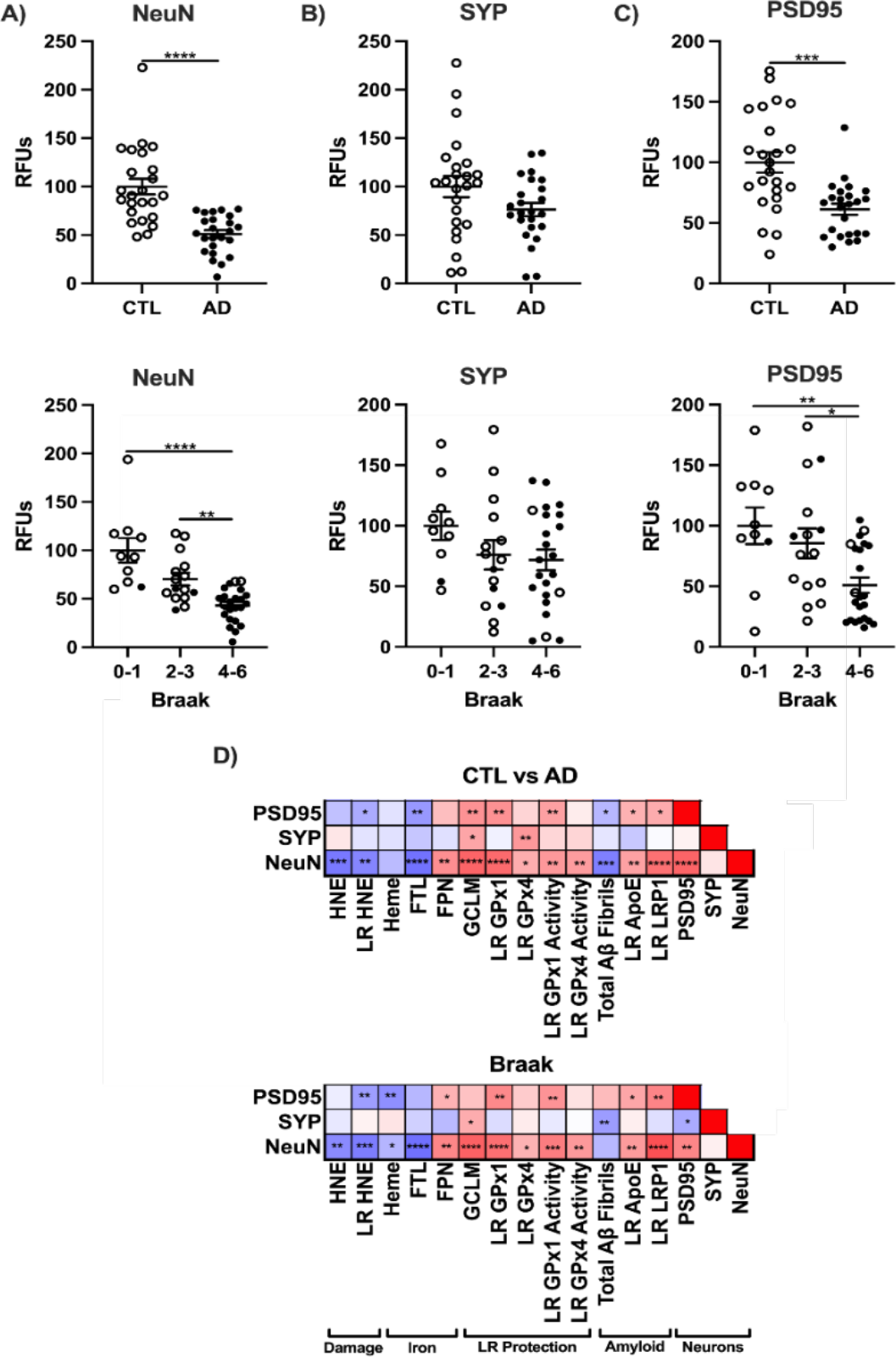
Neuronal loss and transcriptional changes in AD prefrontal cortex. Western blots as relative fluorescent units (RFU) of neuronal markers by CTL vs AD, or Braak stage^41^. **A)** neuronal nuclei antigen (NeuN), **B)** synaptophysin 1 (SYP), **C)** postsynaptic density protein 95 (PSD95). **D)** Correlation matrix for variables with significant relationships to NeuN by CTL vs AD or Braak. Significance, 2-tailed t-test (CTL vs AD), one-way ANOVA (Braak) with Tukey’s posthoc, or Spearman correlation: *p<0.05, **p<0.01, ***p<0.001, ****p<0.0001.

### ApoE allele associations with lipid rafts, lipid peroxidation, and antioxidant defense

Because ApoE4 is associated with greater AD neuropathology, we investigated where our observed changes with iron metabolism, antioxidant defense to lipid rafts, amyloid processing and clearance during AD were associated with ApoE status. Importantly, ApoE4,4 (E4,4) carriers had a significant reduction in NeuN in the cortex, suggesting greater neuronal death is correlated with E4,4 (**Fig. 8A**). Interestingly, cerebellum NeuN was also decreased in E4,4 carriers despite its reported resistance to AD (**Extended Data Fig. E8A**). In addition, we observed a significant decrease in lipid raft cholesterol and proteins in E4,4 cortex (**Fig. 8 B-D**), which suggests that composition of lipid rafts is compromised in E4,4 carriers. As expected, Aβ fibrils were increased in E4,4 cortex, although total LR APP was unaffected (**Fig. 8E-F**). The cerebellum did not display major changes in lipid raft composition across ApoE genotypes (**Extended Data Fig. 8**B-E), although Aβ fibrils were still significantly increased in E4,4, again highlighting that although many markers of pathology are not seen in the cerebellum compared to the cortex, it is not completely devoid of all pathological phenotypes.

**Figure 8:**
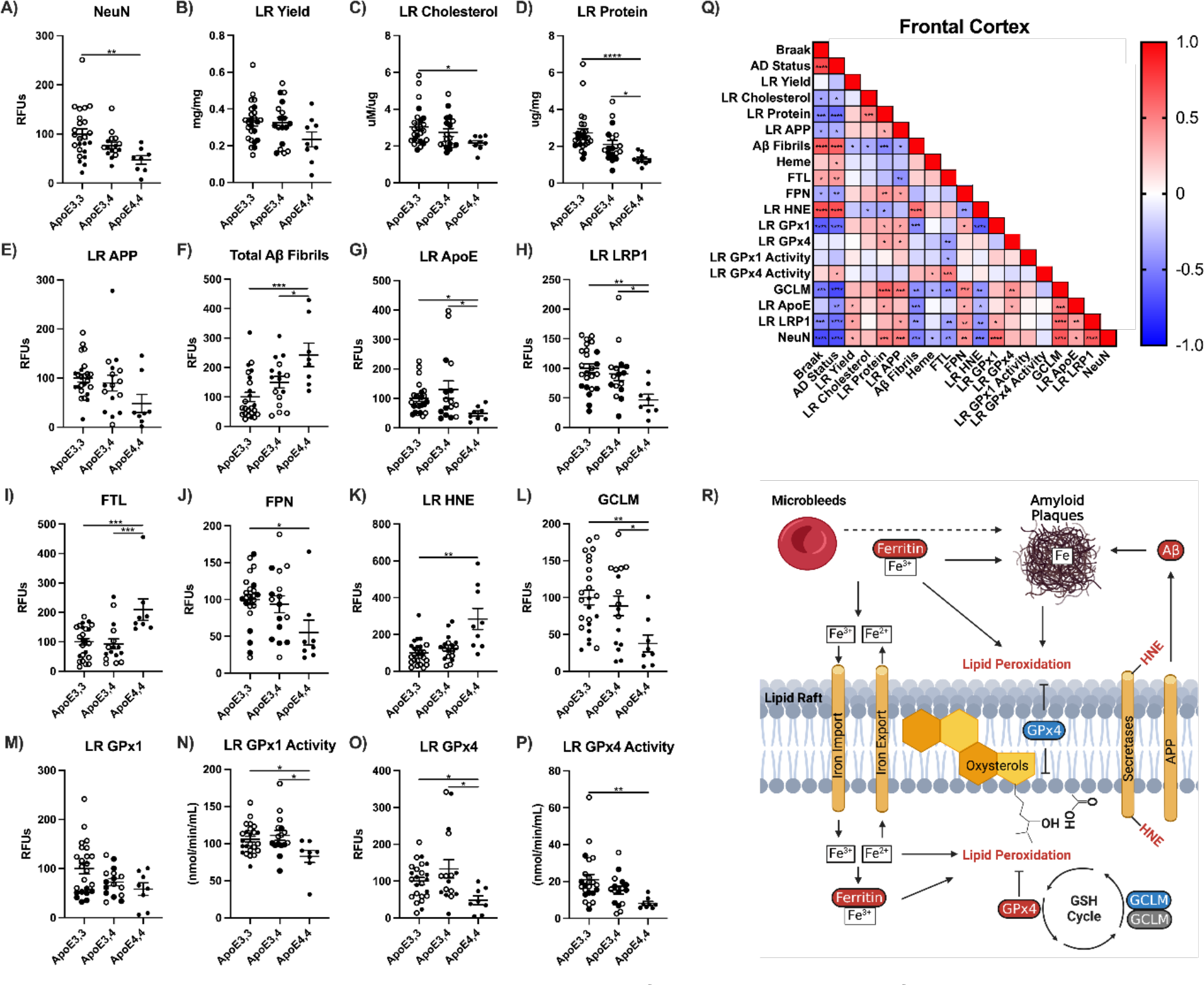
ApoE allele associations with lipid rafts, antioxidant defense, and amyloid in human prefrontal cortex. **A)** NeuN, **B)** LR yield, **C)** LR cholesterol, **D)** total LR protein, **E)** LR APP, **F)** total Aβ fibrils, **G)** LR ApoE, **H)** LR LRP1, **I)** FTL, **J)** FPN, **K)** LR HNE, **L)** GCLM, **M)** LR GPx1, **N)** LR GPx1 activity, **O)** LR GPx4, **P)** LR GPx4 activity. **Q)** Correlation matrix by ApoE allele. One way ANOVA by Tukey’s posthoc, or Spearman correlation: *p<0.05, **p<0.01, ***p<0.001, ****p<0.0001. **R)** Model for ferroptosis in AD.

Next, to determine the consequence of changes in the lipid raft in E4,4 carriers, we measured amyloid clearance proteins ApoE and LRP1 and saw a significant reduction in ApoE4,4 cortex and cerebellum (**Fig. 8G-H, Extended Data** Fig. 8G-H). Interestingly, the iron storage protein FTL was highest in E4,4 cortex with corresponding decreases in the iron export protein FPN (**Fig. 8I,J**).

Importantly, these changes were directly paralleled with lipid peroxidation in the rafts, as HNE adducts increased significantly in E4,4 carriers (**Fig. 8K**). These changes in lipid peroxidation were also observed with corresponding loss of antioxidant capacity, as GCLM, GPx1 activity, and GPx4 levels and activity, although not GPx1 levels, were significantly reduced in E4,4 carriers (**Fig. 8L-P**). Interestingly, although some of these phenotypes were shared across brain regions, most changes were not seen in the cerebellum (**Extended Data Fig. 8**I-P). Overall, our data provide strong evidence that many phenotypes associated with iron-related oxidative damage of lipids, which drive AD pathology, are worse in the E4,4 carriers, consistent with the consensus that E4,4 carriers are at the highest risk for AD. LR markers correlated negatively with Aβ fibrils in cerebral cortex more strongly (**Fig. 8Q**) than cerebellum (**Fig. E8R**). Aβ fibrils also correlated positively with LR HNE levels. The correlation between NeuN and GCLM or FPN remained strong by ApoE allele, as well as for clinical ‘CTL vs AD’ and ‘Braak’ stage. LR GPx1 also positively correlated with NeuN.

## DISCUSSION

Iron is increasingly implicated in brain oxidative damage during AD^21,46,47^. We report the most comprehensive assessment of iron and its associated proteins in postmortem human aging brain. These new data strongly associate iron with antioxidant defenses and lipid peroxidation that vary by brain region for oxidative damage during AD. We introduce an approach to minimize confounds from vascular and entrapped blood by gentle washing that is validated in mouse brain. This increase in oxidative damage of lipids is associated with altered antioxidant defense pathways. Detailed examination of membrane subfractions showed changes specific to lipid rafts where amyloid is processed. ApoE4 allele carriers had increased oxidative damage with AD. The hypothesized role of iron was tested in a mouse model with the iron chelator DFO. In EFAD mice 1-week of DFO treatment solubilized fibrillar amyloid, consistent with the role of iron in amyloid plaques. This release of entrapped iron on amyloid fibrils is relevant to ongoing clinical trials in which monoclonal therapies dissolve amyloid plaques which must release iron.

Heme levels were elevated in AD brains after exogenous blood was minimized. The residual heme may come from leakage from the BBB or microbleeds^48,49^. Heme iron is liberated by heme oxygenase and stored after conversion of ferrous iron to ferric by ferroxidase in FTH1^50^ and then packaged on FTL. Since FTL increased with AD, this is indicative of excess iron in AD despite not observing differences in tissue iron. Plaques and NFTs also contain ferritin protein and iron^46,51–53^ and iron also directly influences amyloid peptide production by the iron-response element in the APP primary transcript^54^. Reexamination of data from Good et al 1992 shows neighboring CA1 neurons without NFTs had 50% less cytoplasmic iron (**Table S2**)^45^. Additionally, ferroportin which is the sole iron export protein was decreased in both brain regions with AD. Lowered FPN exacerbates ferroptosis in neuroblastoma cells treated with erastin^55^. Altogether, these data are consistent with the ferroptotic hypothesis that excess iron is a mechanism in AD pathogenesis.

In addition to changes in iron and iron-related pathways, a loss of detoxification pathways and an increase in oxidative damage contributes to AD pathology. Lipid peroxidation can cause production of reactive alpha-beta unsaturated aldehydes with HNE being the most common. Free HNE is cleared through glutathione s-transferase family members, primarily GSTA4 in the brain, or ALDH2, which is GSH-independent. GSH is needed for most of these processes which has been reported depleted during AD^56,57^, likely from the low levels of GCLM shown in these data. GCLM increases the rate of catalysis over 4-fold for production of γ-glutamylcysteine and is the rate-limiting step in GSH biosynthesis^39^. Cysteine availability is also rate-limiting from its primary source, cystine, which is imported by xCT. Independent of GSH, lipid peroxidation is also attenuated by ferroptosis suppressor protein 1 (FSP1) via the quinol cycle^58^, which we showed is decreased in AD prefrontal cortex and cerebellum. GSH and quinol cycles both rely on NAPDH produced from the pentose phosphate pathway from G6PD and PGD. G6PD is rate-limiting in the formation of NAPDH^59^ and was also decreased in AD prefrontal cortex. All of these antioxidants and iron metabolic proteins are regulated by transcriptional by Nrf2, which is decreased during AD^60,61^, also shown here. Decreases in GCLM, FSP1, and G6PD with AD hinder the antioxidant systems which neutralize lipid peroxidation.

We also introduce the lipid raft to oxidative repair mechanisms in AD. Lipid rafts are tightly packed microdomains rich in cholesterol, sphingomyelin, and phosphatidylcholine within cellular membranes^62,63^. Lipid classes within the raft are easily oxidized enzymatically (lipoxygenases) or nonenzymatically, by Fenton chemistry^64^. Altered composition of lipid classes within lipid rafts impairs biochemical signal transduction^65^ that can cause cell death^66^. Additionally, sterols are more oxidizable that polyunsaturated fatty acids, contrary to prior reports^67^. Oxidation of lipids within rafts may be responsible for the decreased raft yield we reported for AD^41^. ApoE4 carriers also had the lowest raft cholesterol, which introducing new mechanisms for the E4 allele because low cholesterol levels in lipid rafts disrupts raft signaling that can cell death^68^.

Membrane lipids are protected by GPx4 and Prdx6, which chemically reduce oxidized phospholipids and sterols with GSH^69,70^. We defined lipid rafts for glutathione peroxidases (GPx1-8) which contained only GPx1 and GPx4 but not Prdx6. Prdx6 and GPx1 also reduce lipid hydroperoxides^70^ released from membrane lipids^71^ and may therefore play a secondary role in defense against ferroptosis. The absence of Prdx6 in the lipid raft fraction is consistent with its transient interactions with membranes^72,73^. AD rafts had lower GPx4 and GPx1 protein and correspondingly lower enzyme activity, consistent with the prominent lipid peroxidation in AD. The non-raft membrane fraction showed minimal oxidation with AD. Furthermore, baseline GPx4 is 2-fold higher in lipid rafts of normal brains than in the non-raft membrane fraction, suggesting that lipid rafts are hotspots for iron-mediated oxidative damage. The observed increases in whole cell GPx4, but with decreased lipid raft GPx4, suggests a localization issue. Cytosolic GPx4 is 20 kDa, while the mitochondrial isoform of 23 kDa containing a leader sequence cleaved for mitochondrial membrane localization^74^. It is possible leader sequences may be needed for membrane localization. Localization may also be mediated by a phosphorylation site that localizes GPx4 to membranes^75^.

Because lipid rafts are the main site of amyloid production, their lipid peroxidation may alter APP processing enzymes. APP processing is enhanced by the γ-secretase in rat cortex neurons and human neuronal cells exposed to 1-10 µM HNE or 100 µM FeSO4^76^. Moreover, H2O2 increased BACE1 activity, the primary secretase responsible for amyloid peptide production in mouse neuronal N2a cells transgenic for human APP695^77^. HNE adducts can impact enzyme activities including GCLM, Prdx6, G6PD, among others relevant to antioxidant defense^78^. APP processing enzymes contain at least 10% of the amino acids His, Lys, Arg, and Tyr that can be covalently linked by HNE or nitrotyrosine. BACE1 (57/501), PSEN1 (50/467), ADAM10 (142/748). BACE1 activity also increases with AD in prefrontal cortex^79^ and serum^80,81^ paralleling the increases of lipid peroxidation in serum and tissue^82,83^. Future studies will evaluate how lipid raft oxidation alters APP processing for amyloid peptides.

AD also decreased Nrf2, a transcriptional regulator of gene responses for oxidative repair that include glutathione peroxidases and GSH biosynthesis proteins^84,85^. Besides our data showing Nrf2 levels decline by 75% in the AD cortex but not cerebellum, previous work from coauthors showed decline of Nrf2 mediated antioxidant defenses in aging mouse cerebellum^86,87^. We suggest that the age-related decline of Nrf2 defense enhances membrane oxidation that may increase amyloid peptide production from microbleed iron. There were few changes in iron-related pathways, despite decreased expression of Nrf2 related genes. However, CD163, the hemoglobin-haptoglobin scavenger which increases in response to microbleeds, was shared between brain regions with AD and was among the top responding mRNAs^44^. CD163 is found in microglia^88^, which have high levels of ferritin in AD that may arise from microbleeds^53,89,90^. Endogenous cell ferritin merits consideration because microglia and astrocytes in young rat brain have 2-fold more ferritin than neurons, while oligodendroglia have the highest ferritin levels^91^.

New AD changes are shown for multiple transcription factors. While Nrf2 is greatly lowered in AD (see above), there are major increases of NF-kB and NCOA4. NF-kB regulates several pro-oxidant enzymes while NCOA4 responds to iron excess for by lysosomal degradation iron bound proteins (ferritinophagy)^43^. A relationship of NF-kB to membrane oxidation is suggested by its role in ferroptosis of glioblastoma cells treated with a GPx4 inhibitor^92^. Future studies will supersede this bulk analysis which cannot resolve cell type specificity for iron-mediated damage.

Finally, ApoE4,4 carriers had the greatest oxidative damage in AD brains and lowest antioxidant protection in lipid rafts, which contrasts with an *in vitro* study of rat neurons with human ApoE alleles^93^. ApoE alleles may also differentially alter antioxidant gene regulation in brain consistent with their roles in Aβ clearance and inflammation^94^. Our findings introduce the ApoE alleles as important to iron-mediated processes for further discussion of ferroptosis.

Lastly, we experimentally tested the role of iron with acute deferoxamine (DFO) in EFAD mice. Chelation therapy for AD was first attempted in 1991 to remove aluminum by DFO, which slowed cognitive decline during 24-months^95^. While the cognitive benefits of DFO were attributed to aluminum chelation, Gleason and Bush^96^ also noted that DFO has 6-fold greater affinity for ferric iron than aluminum. Our DFO data directly supports the major role of iron in multiple AD pathologies, as DFO reduced HNE oxidation and decreased amyloid fibrils. This suggests that iron chelation may solubilize amyloid fibrils to monomers while liberating iron. Traumatic brain injury, which often results in microbleeds, shows increases in amyloid levels that are attenuated with DFO treatment^97^. We previously showed that microbleeds precede amyloid formation and that these two pathologies colocalize in the EFAD mice^98^. Is it possible that amyloid production has a role in preventing more heme-derived iron from entering and depositing in the brain parenchyma? Furthermore, if iron is liberated from amyloid fibrils, then oxidative damage would be expected in the absence of chelation. DFO treatment increased Nrf2 and downstream antioxidants together with decreased oxidative damage, that support DFO clinical value. Current amyloid monoclonal therapies that target aggregated amyloid must also liberate labile iron. The liberated iron could contribute to amyloid related imaging abnormalities (ARIA) of clinical trials^99–101^. Combination therapies should be considered for iron chelators with amyloid monoclonal treatment.

In summary, this analysis of iron metabolism and membrane oxidation in AD is the first to comprehensively document the network of proteins that mediate iron transport and GSH-mediated neuroprotection relevant to ferroptosis. AD pathway specificity was demonstrated by the greater changes in the prefrontal cortex than cerebellum, consistent with quantitative differences in their AD neuropathology. Our data identify novel alterations in iron related transport, clearance, and oxidative repair mechanisms in relation to lipid rafts and APP processing. Moreover, multiple anti-ferroptotic pathways are implicated by the decreases in GCLM, FSP1, FPN, and lipid raft GPx4. Experimental reduction of lipid peroxidation by iron chelation may be considered for co-therapy with monoclonal antibody treatments to reduce amyloid that inevitably releases iron associated with amyloid plaques. Taken together, this suite of findings confirms that iron can contribute to AD pathology by exacerbating oxidative damage.

## METHODS

### Human Samples

Postmortem prefrontal cortex and cerebellum with Braak scores were provided by three Alzheimer’s Disease Centers: University of Southern California (Dr. Carol Miller), University of California Irvine (UCI MIND), and University of Washington (BRaIN). Samples were genotyped for ApoE alleles by PCR for SNP variants rs429358 and rs7412. All human subjects provided informed consent. Human tissue use was approved through IRB protocol #UP-20-00014-EXEMPT. Details for individual brains are in **Supplementary Table 1**.

### Mice

C57BL/6 mice from Jackson laboratories or ApoEFAD mice from Mary Jo LaDu (University of Illinois, Chicago) were housed by the University of Southern California’s Department of Animal Resources. Mice had ad libitum access to Purina Lab Chow (LabDiet, Hayward, CA) and water. Mice were housed in groups of 5 at 22°C/ 30% humidity, with standard nesting material and light cycles of 0600-1800 hr. Studies were approved by the USC Animal Use and Care Committee (IACUC# 20417). Female ApoEFAD mice were given DFO treatment either in rodent chow (10mg/kg/day) for two weeks or by intraperitoneal injection twice daily for one week (10m/kg/day). At 6-months old mice were anesthetized with isoflurane and euthanized by cardiac puncture. C57BL/6 mouse cortex was collected as non-perfused, with transcardial PBS perfusion, or PBS perfusion + 10U/mL heparin (Sigma-Aldrich, St. Louis, MO). Tissues were stored at -80°C.

### Tissue Washing

Human and mouse brain tissue was thawed on ice and minced by ceramic scalpel for iron and heme measurements or by surgical scissors. PBS (10 volumes) was placed in each tube and the samples were gently washed by two cycles of vortexing and collected by gentle centrifugation 800g/30 sec/4°C. PBS was retained on a small subset of samples and further spun at 4,000g/10 min/ 4°C. The pellet was sonicated 10 sec at 50% power in PBS. Resuspended pellet and lysate were assayed for heme (**Table 1; Fig. S1**).

### Amyloid Measurements

Human tissue was processed for amyloid measurements as previously described^41^. Briefly, tissue was homogenized in RIPA without SDS and spun for 10,000g for 1h at 4°C. Supernatant was collected and analyzed for soluble Aβ fibrils using M78 (Conformation specific antibody for Aβ fibrils^102^, a gift from Dr. Charles Glabe, UCI). The residual pellet was resuspended in 70% formic acid and sonicated. The resuspension was nutated at room temperature for 2h and neutralized with 20 volumes of 1M Tris and concentrated with a vacuum concentrator (Labconco, Kansas City, MO). Amyloid levels were assayed by dot blot and visualized as described below for Western blots.

### Lipid Raft and Non-Raft Membrane

The lipid raft fraction was isolated from 40 mg tissue homogenates of frozen prefrontal cortex (Brodmann area 8, 9 or 10) and cerebellum using a column-based isolation kit (Invent Biotechnologies, Plymouth, MN). These findings were verified by sucrose gradient ultracentrifugation^103^. Lipid raft and the remaining detergent soluble non-raft membrane (NRM) fractions were resuspended in PBS+1% triton x-100 and sonicated 10 seconds at 50% power. Protein was assayed by the copper-based 660 nm assay (ThermoFisher Scientific, Waltham, MA).

Lipid raft enrichment and purity were determined by cytosolic GAPDH, several lipid raft markers including flotillin1, and nuclear marker H3^41^.

### Nuclear Isolation

20 mg of frozen brain was homogenized in 125uL of sucrose buffer for nuclear and cytosolic fractions. A previously published protocol^104^ was modified for brain tissue by adding 1% NP- 40 to the first wash step, resuspending the pellet, and incubating on ice for 30 min. Nuclear and cytosolic extractions were cross probed for purity against both H3 and GAPDH by Western blot^105^ and showed the expected enrichment (**Fig. S5**).

### Western Blot

Proteins from lipid rafts (5 ug), whole cell lysate (20 ug), or nuclear fractions (20 ug) were heated at 75°C under denaturing conditions and resolved on 4-20% gradient gels. Proteins were electroblotted using a Criterion blotter (Bio-Rad Laboratories, Hercules, CA) and transferred onto 0.45um polyvinyl difluoride membranes. Membranes were stained using Revert 700 fluorescent protein stain and imaged prior to blocking with LI-COR Intercept blocking buffer (LI-COR Biosciences, Lincoln, NE), followed by primary antibodies. Membranes were incubated with IRDye 800CW and/or 700CW secondary antibodies and visualized by Odyssey (LI-COR Biosciences). Western blot data was quantified with ImageJ and normalized by total protein per lane and/or loading control protein.

### Biochemical Assays

Lipid rafts were assayed for total cholesterol (Cell Biolabs, San Diego, CA). Tissue hemoglobin or heme quantified by Quantichrom assay (BioAssay Systems, Hayward, CA). α- Hemoglobin was quantified by ELISA (ThermoFisher Scientific, Waltham, MA). Total glutathione peroxidase activities were measured by enzymatic assay (Cayman Chemical, Ann Arbor, MI). GPx4 activity was assayed by generating oxidized phosphatidylcholine using soybean lipoxygenase^106^. Purified PCOOH was used in place of cumene hydroperoxide in Cayman’s total GPx activity assay at a concentration of 30µM per well.

### ICP-MS

50mg of brain tissue was cut with a ceramic scalpel and placed in metal free certified test tubes. Tissues were washed twice with PBS and homogenized in PBS by polyethylene pestle (PES- 15-B-SI, Axygen) with purified water (18.2 MΩ; Millipore, Burlington, MA) treated with Chelex 100 (sigma, St. Louis, MO). Homogenates were desiccated by vacuum centrifuge at 95°C for 90 min. and solubilized in trace metal free 70% HNO3 overnight. 30% H2O2 was then added followed by dessication. After resuspension in 2% HNO3, samples were analyzed by Agilent 7500ce ICP-MS in hydrogen mode with a practical detection limit of 10ppb and a relative standard deviation (RSD) of replicate measures between 0.2 to 5%. Total iron concentration was normalized to wet weight tissue.

### RNA-Sequencing

20mg of brain tissue was homogenized in TRIzol reagent using a BeadBug Benchtop Homogenizer. Brain tissue was suspended in 1 mL of TRIzol and homogenized for 6 rounds of 10 s homogenization and 60 s holds between each round. 300 µL of chloroform was added to the sample and aqueous separation of RNA was performed using centrifugation in a heavy gel phase-lock tube (VWR, 10847-802). The aqueous phase was applied to a standard column-based RNA purification kit (Quantabio, Extracta Plus, Cat# 95214-050) following the manufacturer’s protocol. Library preparation and RNA sequencing was performed by Novogene. Reads from samples with were processed using trim_galore (0.6.5-1) and mapped to the GRCh38.p14 genome with STAR-2.7.10a. Mapped reads were counted to genes using featureCounts (Subread-2.0.3) and the GRCh38.108 GTF file. Unwanted noise and artifacts were removed using svaseq and removeBatchEffects from the sva package-3.42.0 and differential expression analysis was performed with DESeq2-1.34.0 using R-4.3.2. Raw RNA-seq data is available through Annotare: E-MTAB- 14167.

### Statistics

GraphPad Prism 9 software (GraphPad, San Diego, CA) was used for graphing and statistical analysis. Several measures were adjusted by age and sex by multiple linear regression which yielded no significant interactions. For these reasons data were analyzed by two-tailed t-test or one-way ANOVA with Tukey’s HSD (p<0.05) with a confidence interval set at 95%. Data that were not normally distributed were analyzed by Mann Whitney U or Kruskal-Wallis with Dunn’s test. All correlation matrixes used Spearman’s correlation for non-linear data.

## Acknowledgements & Funding

The authors are grateful to Liying Zhao for performing ICP-MS measurements, and Carol Church and Kymry Jones for assistance with postmortem tissues. The authors appreciate the helpful discussions of brain samples with Carol Miller (USC) and C. Dirk Keene (UW). Tissue for this study was obtained from the USC ADRC Neuropathology Core, NIAAG066530. The UCI-ADRC is funded by NIH/NIA Grant P30AG066519. J. Silva and E. Head were supported by P30AG066519. Lab studies were supported by NIH grants to CEF (R01-AG051521, P50-AG05142, P01-AG055367) and Cure Alzheimer’s Fund; Larry L. Hillblom Foundation Grant 2022-A-010-SUP, the Glenn Foundation for Medical Research and AFAR Grant for Junior Faculty Award, and Navigage Foundation Award to RHS; T32AG052374 to GG; and T32-AG000037 (PI: Eileen Crimmins) to MAT. Brain specimens were obtained from ADRC Tissue Cores: USC (P50-AG005142, AG066530); University of California Irvine (P30-AG066519); UW (P30-AG066509; U01-AG006781). Venn diagrams were made with Venny 2.1.0 and schematics made using BioRender.

## Ethics Declarations

The authors declare no competing interests.

## Data Availability

The authors declare that the data supporting the findings of this study are available within the paper and its supplementary information files. Should any raw data files be needed in another format they are available from the corresponding author upon reasonable request.

---

**Supplemental Figure 1:**
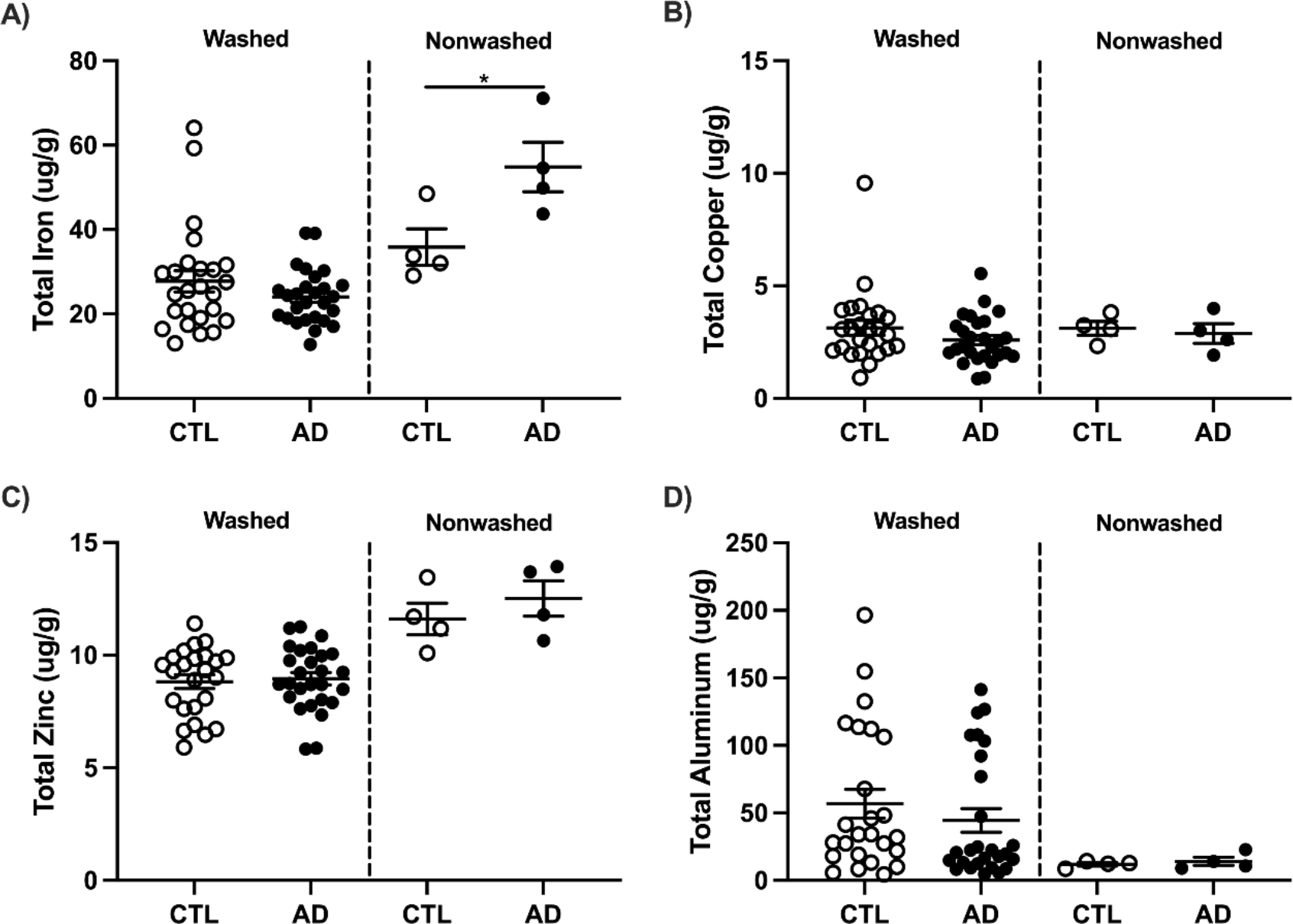
Total **A)** Iron, **B)** Copper, **C)** Zinc, and **D)** Aluminum concentrations by ICP-MS (Table 1). **A-D,** Human prefrontal cortex was washed with PBS or non-washed. Significance by 2-tailed t-test (CTL vs AD): *p<0.05.

**Supplemental Figure 2:**
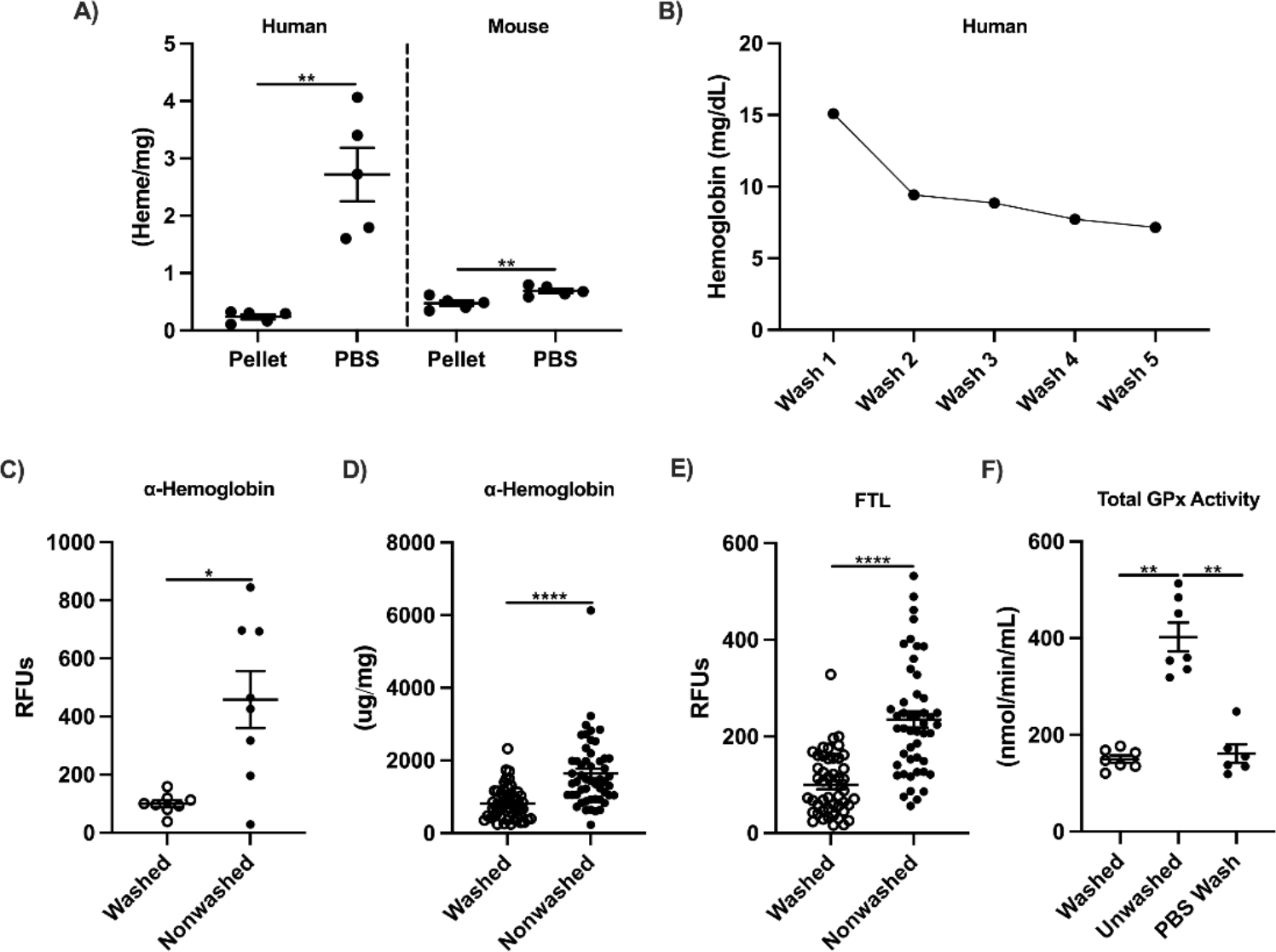
Blood on tissues influences iron and antioxidant measurements. **A)** Nonperfused mouse and washed human cerebral cortex. **B)** Collected wash was assayed for heme in 5 consecutive washes; washes 1 and 2 reduced makers by 90%; washes 3-5 yielded <10% further decreases. The lower α- hemoglobin in washed human frontal cortex was confirmed by **C)** Western blot, **D)** immunoassay, **E)** Western blot for FTL, and **F)** total GPx activity. Significance, 2-tailed t- test (A, C-E) or one-way ANOVA with Tukey’s post-hoc (F): *p<0.05, **p<0.01, ***p<0.001, ****p<0.0001.

**Supplemental Figure 3:**
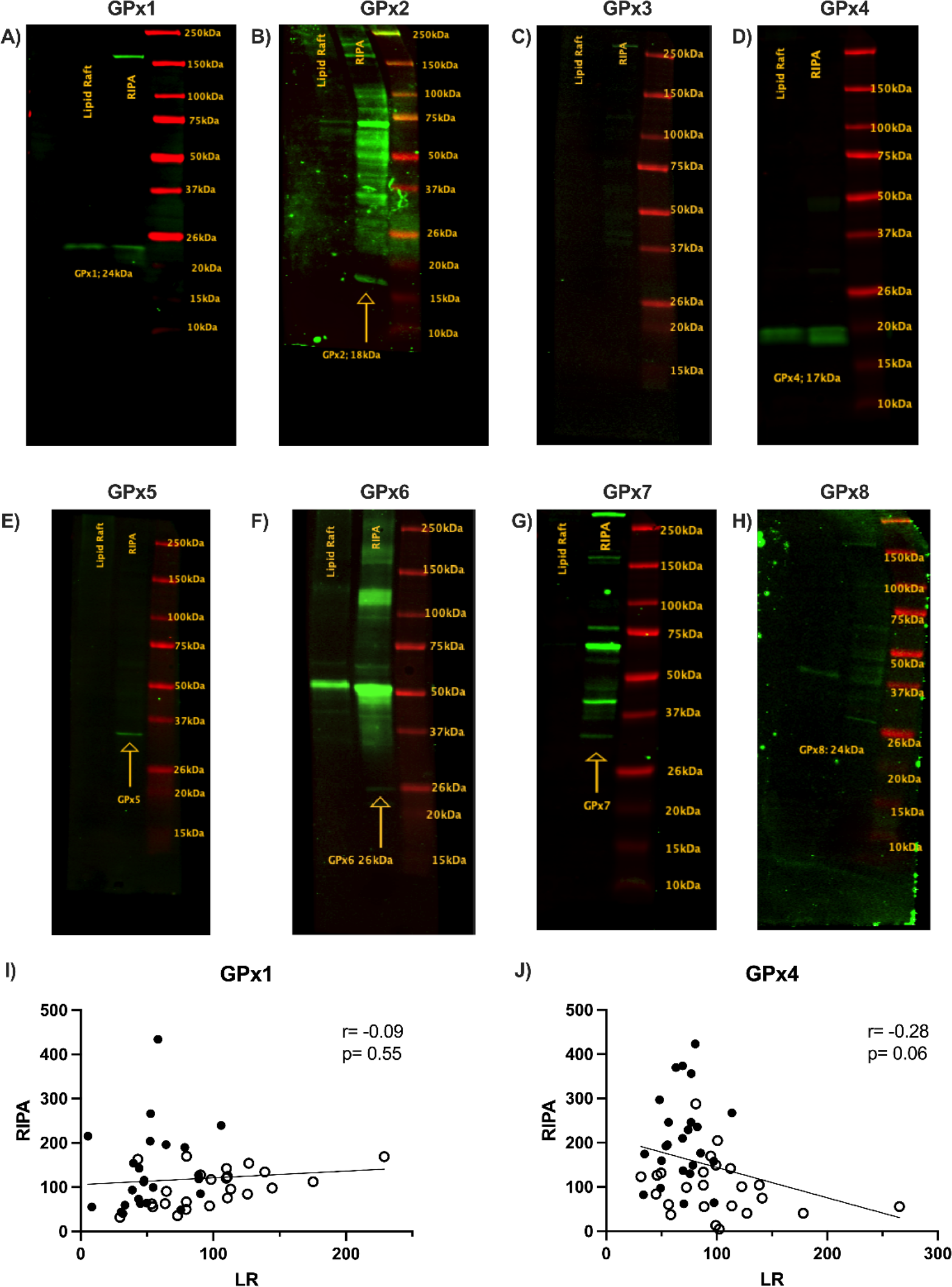
Glutathione peroxidases in lipid rafts from human prefrontal cortex. The GPx isoforms were resolved immunologically on Western blots of lipid raft and whole cell lysates (RIPA buffer extract). **A)** GPx1, **B)** GPx2, **C)** GPx3, **D)** GPx4, **E)** GPx5, **F)** GPx6, **G)** GPx7, **H)** GPx8. Coplots of **I)** GPx1 and **J)** GPx4 for RIPA extract and lipid raft fractions.

**Supplemental Figure 4:**
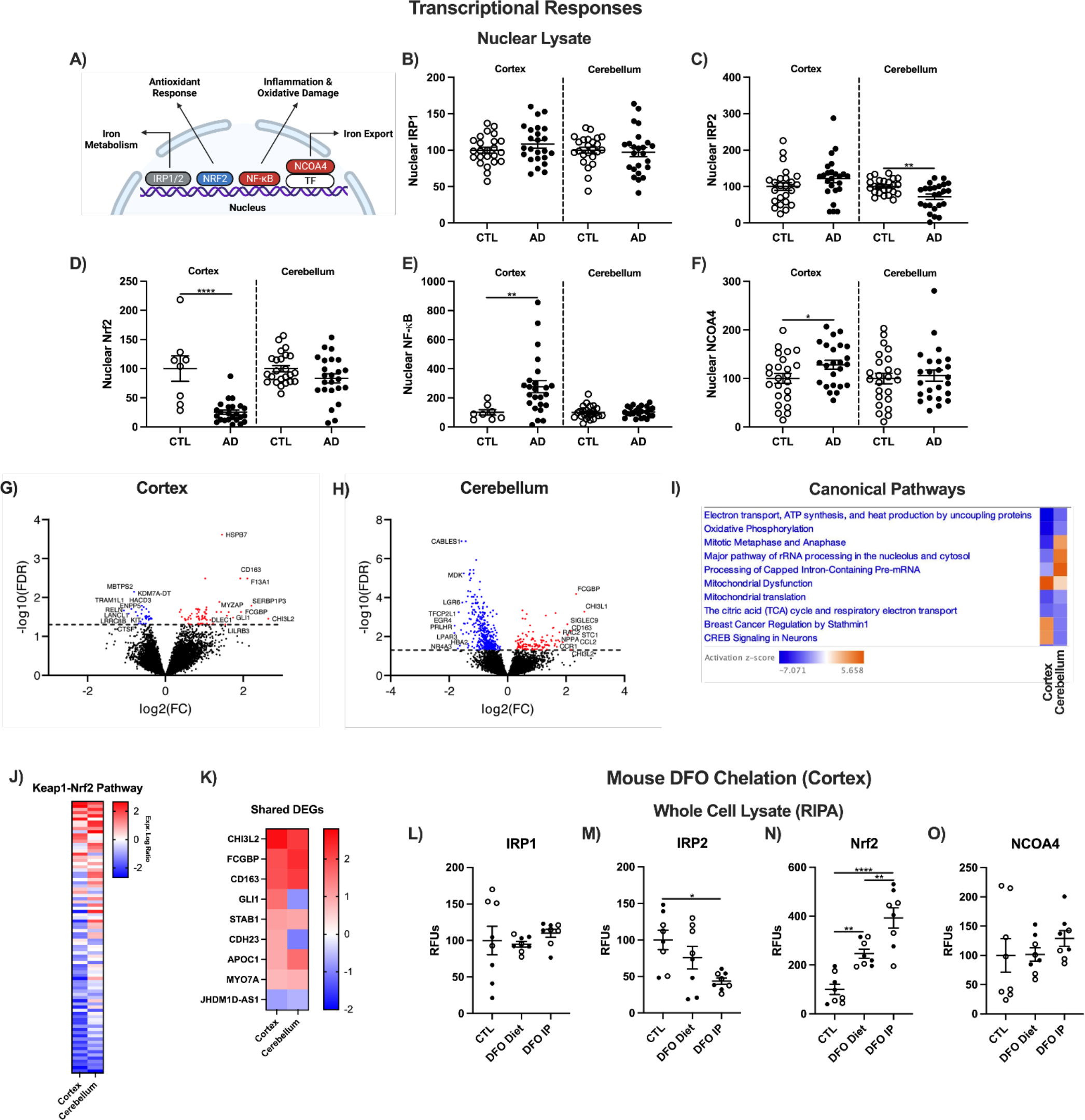
Transcriptional responses mediating iron homeostasis. **A)** Transcription factors of genes for antioxidant defense and iron signaling in human prefrontal cortex of AD and CTL. IRP, iron regulatory protein; Nrf2, nuclear factor erythroid 2–related factor 2; NF-κB p65, nuclear factor κ-light-chain-enhancer of activated B cells; NCOA4, nuclear receptor coactivator 4; TF, transcription factor. Western blot data from nuclear lysates as relative fluorescent units (RFUs) for **B)** IRP1, **C)** IRP2, **D)** Nrf2, **E)** NF-κB p65, and **F)** NCOA4 in prefrontal cortex and cerebellum. Volcano plots of DEGs from RNA-seq for **G)** prefrontal cortex, **H)** cerebellum, and **I)** top shared canonical pathways. **J)** Keap1- Nrf2 pathway and **K)** shared DEGs between cortex and cerebellum. DFO treated EFAD mouse cortex total (RIPA lysate) for **L)** IRP1, **M)** IRP2, **N)** Nrf2, and **O)** NCOA4 protein in. Significance by 2-tailed t-test (B-F) or One way ANOVA with Tukey’s posthoc test (L-O): *p<0.05, **p<0.01, ****p<0.0001.

**Supplemental Figure 5:**
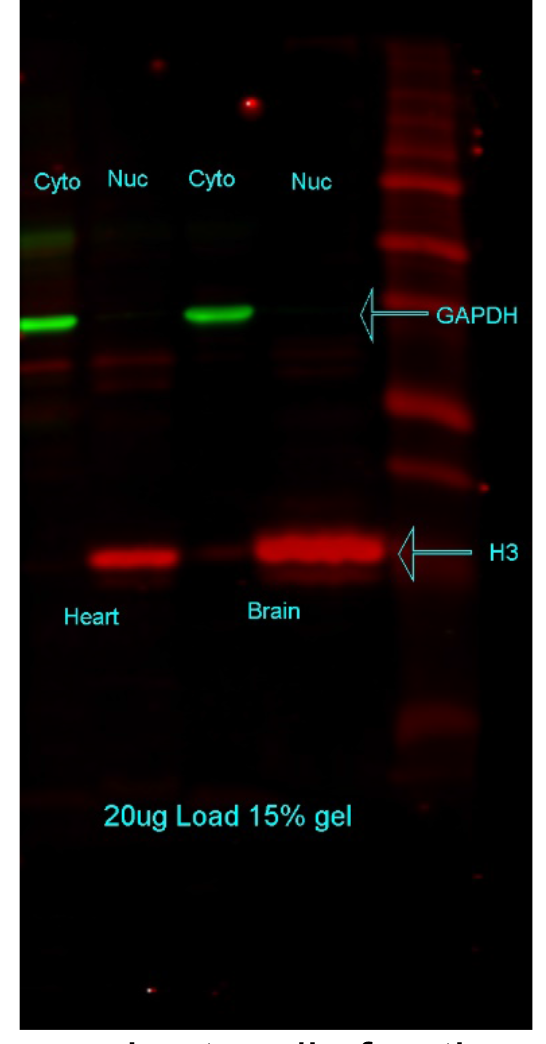
Nuclear and cytosolic fractions from mouse heart and cortex. Lysates probed by Western blot for histone 3 (H3, nuclear marker) and glyceraldehyde 3-phosphate dehydrogenase (GAPDH, cytosolic marker).

**Supplemental Table 1:**
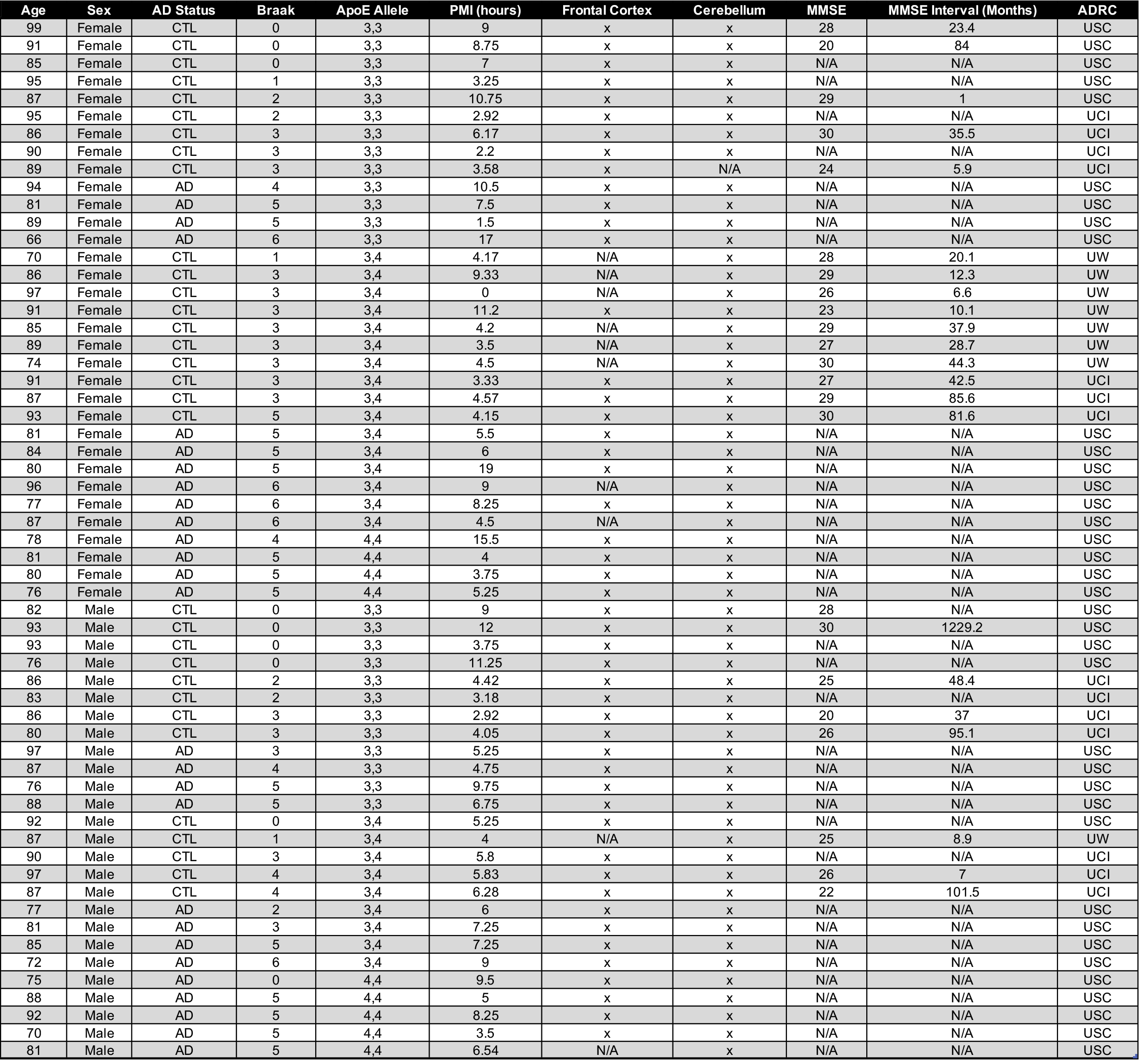
Data shown for brain specimens by each ADRC source: USC, 37; UCI, 14; UW, 8. White, 76%; Latino, 14%; African American, 3%; Asian, 2%; N/A, 5%. ApoE3,3, 42%, ApoE3,4, 42%, ApoE4,4, 16%.

| Age | Sex | AD Status | Braak | ApoE Allele | PMI (hours) | Frontal Cortex | Cerebellum | MMSE | MMSE Interval (Months) | ADRC |
| --- | --- | --- | --- | --- | --- | --- | --- | --- | --- | --- |
| 99 | Female | CTL | 0 | 3,3 | 9 | x | x | 28 | 23.4 | USC |
| 91 | Female | CTL | 0 | 3,3 | 8.75 | x | x | 20 | 84 | USC |
| 85 | Female | CTL | 0 | 3,3 | 7 | x | x | N/A | N/A | USC |
| 95 | Female | CTL | 1 | 3,3 | 3.25 | x | x | N/A | N/A | USC |
| 87 | Female | CTL | 2 | 3,3 | 10.75 | x | x | 29 | 1 | USC |
| 95 | Female | CTL | 2 | 3,3 | 2.92 | x | x | N/A | N/A | UCI |
| 86 | Female | CTL | 3 | 3,3 | 6.17 | x | x | 30 | 35.5 | UCI |
| 90 | Female | CTL | 3 | 3,3 | 2.2 | x | x | N/A | N/A | UCI |
| 89 | Female | CTL | 3 | 3,3 | 3.58 | x | N/A | 24 | 5.9 | UCI |
| 94 | Female | AD | 4 | 3,3 | 10.5 | x | x | N/A | N/A | USC |
| 81 | Female | AD | 5 | 3,3 | 7.5 | x | x | N/A | N/A | USC |
| 89 | Female | AD | 5 | 3,3 | 1.5 | x | x | N/A | N/A | USC |
| 66 | Female | AD | 6 | 3,3 | 17 | x | x | N/A | N/A | USC |
| 70 | Female | CTL | 1 | 3,4 | 4.17 | N/A | x | 28 | 20.1 | UW |
| 86 | Female | CTL | 3 | 3,4 | 9.33 | N/A | x | 29 | 12.3 | UW |
| 97 | Female | CTL | 3 | 3,4 | 0 | N/A | x | 26 | 6.6 | UW |
| 91 | Female | CTL | 3 | 3,4 | 11.2 | x | x | 23 | 10.1 | UW |
| 85 | Female | CTL | 3 | 3,4 | 4.2 | N/A | x | 29 | 37.9 | UW |
| 89 | Female | CTL | 3 | 3,4 | 3.5 | N/A | x | 27 | 28.7 | UW |
| 74 | Female | CTL | 3 | 3,4 | 4.5 | N/A | x | 30 | 44.3 | UW |
| 91 | Female | CTL | 3 | 3,4 | 3.33 | x | x | 27 | 42.5 | UCI |
| 87 | Female | CTL | 3 | 3,4 | 4.57 | x | x | 29 | 85.6 | UCI |
| 93 | Female | CTL | 5 | 3,4 | 4.15 | x | x | 30 | 81.6 | UCI |
| 81 | Female | AD | 5 | 3,4 | 5.5 | x | x | N/A | N/A | USC |
| 84 | Female | AD | 5 | 3,4 | 6 | x | x | N/A | N/A | USC |
| 80 | Female | AD | 5 | 3,4 | 19 | x | x | N/A | N/A | USC |
| 96 | Female | AD | 6 | 3,4 | 9 | N/A | x | N/A | N/A | USC |
| 77 | Female | AD | 6 | 3,4 | 8.25 | x | x | N/A | N/A | USC |
| 87 | Female | AD | 6 | 3,4 | 4.5 | N/A | x | N/A | N/A | USC |
| 78 | Female | AD | 4 | 4,4 | 15.5 | x | x | N/A | N/A | USC |
| 81 | Female | AD | 5 | 4,4 | 4 | x | x | N/A | N/A | USC |
| 80 | Female | AD | 5 | 4,4 | 3.75 | x | x | N/A | N/A | USC |
| 76 | Female | AD | 5 | 4,4 | 5.25 | x | x | N/A | N/A | USC |
| 82 | Male | CTL | 0 | 3,3 | 9 | x | x | 28 | N/A | USC |
| 93 | Male | CTL | 0 | 3,3 | 12 | x | x | 30 | 1229.2 | USC |
| 93 | Male | CTL | 0 | 3,3 | 3.75 | x | x | N/A | N/A | USC |
| 76 | Male | CTL | 0 | 3,3 | 11.25 | x | x | N/A | N/A | USC |
| 86 | Male | CTL | 2 | 3,3 | 4.42 | x | x | 25 | 48.4 | UCI |
| 83 | Male | CTL | 2 | 3,3 | 3.18 | x | x | N/A | N/A | UCI |
| 86 | Male | CTL | 3 | 3,3 | 2.92 | x | x | 20 | 37 | UCI |
| 80 | Male | CTL | 3 | 3,3 | 4.05 | x | x | 26 | 95.1 | UCI |
| 97 | Male | AD | 3 | 3,3 | 5.25 | x | x | N/A | N/A | USC |
| 87 | Male | AD | 4 | 3,3 | 4.75 | x | x | N/A | N/A | USC |
| 76 | Male | AD | 5 | 3,3 | 9.75 | x | x | N/A | N/A | USC |
| 88 | Male | AD | 5 | 3,3 | 6.75 | x | x | N/A | N/A | USC |
| 92 | Male | CTL | 0 | 3,4 | 5.25 | x | x | N/A | N/A | USC |
| 87 | Male | CTL | 1 | 3,4 | 4 | N/A | x | 25 | 8.9 | UW |
| 90 | Male | CTL | 3 | 3,4 | 5.8 | x | x | N/A | N/A | UCI |
| 97 | Male | CTL | 4 | 3,4 | 5.83 | x | x | 26 | 7 | UCI |
| 87 | Male | CTL | 4 | 3,4 | 6.28 | x | x | 22 | 101.5 | UCI |
| 77 | Male | AD | 2 | 3,4 | 6 | x | x | N/A | N/A | USC |
| 81 | Male | AD | 3 | 3,4 | 7.25 | x | x | N/A | N/A | USC |
| 85 | Male | AD | 5 | 3,4 | 7.25 | x | x | N/A | N/A | USC |
| 72 | Male | AD | 6 | 3,4 | 9 | x | x | N/A | N/A | USC |
| 75 | Male | AD | 0 | 4,4 | 9.5 | x | x | N/A | N/A | USC |
| 88 | Male | AD | 5 | 4,4 | 5 | x | x | N/A | N/A | USC |
| 92 | Male | AD | 5 | 4,4 | 8.25 | x | x | N/A | N/A | USC |
| 70 | Male | AD | 5 | 4,4 | 3.5 | x | x | N/A | N/A | USC |
| 81 | Male | AD | 5 | 4,4 | 6.54 | N/A | x | N/A | N/A | USC |

**Supplemental Table 2:** Iron in NFT free and bearing neurons presented as mass charge ratio (m/z) by Good et al. 1992^45^. Significance analyzed in this report by 2-tailed t-test: *p<0.05.

|  | NFT-Bearing Neurons |  |  |  | NFT-Free Neurons |  |  |
| --- | --- | --- | --- | --- | --- | --- | --- |
|  | NFT | Cytoplasm | Nucleus | Neuropil | Cytoplasm | Nucleus | Neuropil |
| **AD** | 65 ± 9 | 20 ± 3 | 42 ± 5 | 14 ± 3 | 12 ± 2* | 44 ± 7* | 11 ± 1* |
| **CTL** | N/A | N/A | N/A | N/A | 4 ± 0.5 | 17 ± 2 | 4 ± 0.5 |

**Supplemental Table 3:** AD change in prefrontal cortex (Brodmann 8-10) and cerebellum by ApoE3 and 4 alleles. FTL, ferritin light chain; FSP1, ferroptosis suppressor protein 1; GPx4, glutathione peroxidase 4. t-test *p<0.05, **p<0.01, ***p<0.001, ****p<0.0001; n.s, not significant.

| Measurement | Cortex<br>(AD vs<br>CTL) | Cerebellum<br>(AD vs<br>CTL) | Cortex<br>(ApoE3,3 vs<br>3,4 & 4,4) | Cerebellum<br>(ApoE3,3 vs<br>3,4 & 4,4) | Cortex<br>Male<br>(AD vs<br>CTL) | Cortex<br>Female<br>(AD vs<br>CTL) |
| --- | --- | --- | --- | --- | --- | --- |
| **FTL** | +65% ** | +10%, n.s. | +30%, n.s. | -30%, n.s. | +55%,<br>n.s. | +75% * |
| **FSP1** | -40% **** | -30% **** | -40% * | -20% ** | -50%,<br>n.s. | -30% * |
| **GCLM** | -60% **** | -35% * | -30%, n.s. | -15%, n.s. | -65% **** | -55% *** |
| **GPx4** | +110%<br>**** | -15% * | +45% * | -30, n.s. | +45%,<br>n.s. | +210%<br>*** |
| **GPx4, LR** | -30% ** | -40% **** | -5%, n.s. | -5%, n.s. | -25% ** | -35%,<br>n.s. |
| **GPx4 Activity,**<br>**LR** | -40% * | -10%, n.s. | -50% ** | -15%, n.s. | -40%,<br>n.s. | -40%,<br>n.s. |
| **Heme** | +25% * | +30% ** | +5%, n.s. | +10%, n.s. | +20%,<br>n.s. | +25%,<br>n.s. |
| **HNE, Total** | +60%<br>**** | -20%, n.s. | +35% ** | -25% * | +35% ** | +90% *** |
| **HNE, LR** | +160%<br>**** | +45% ** | +80% ** | +35% * | +170% ** | +180%<br>*** |
| **NF-κB, nuclear** | +175% ** | +5%, n.s. | +70%, n.s. | +5%, n.s. | +45%,<br>n.s. | +290% ** |
| **Nrf2, nuclear** | -75% **** | -15%, n.s. | -65% *** | -15%, n.s. | -85% ** | -60% ** |

## Extended Figures

**Extended data Figure 1:**
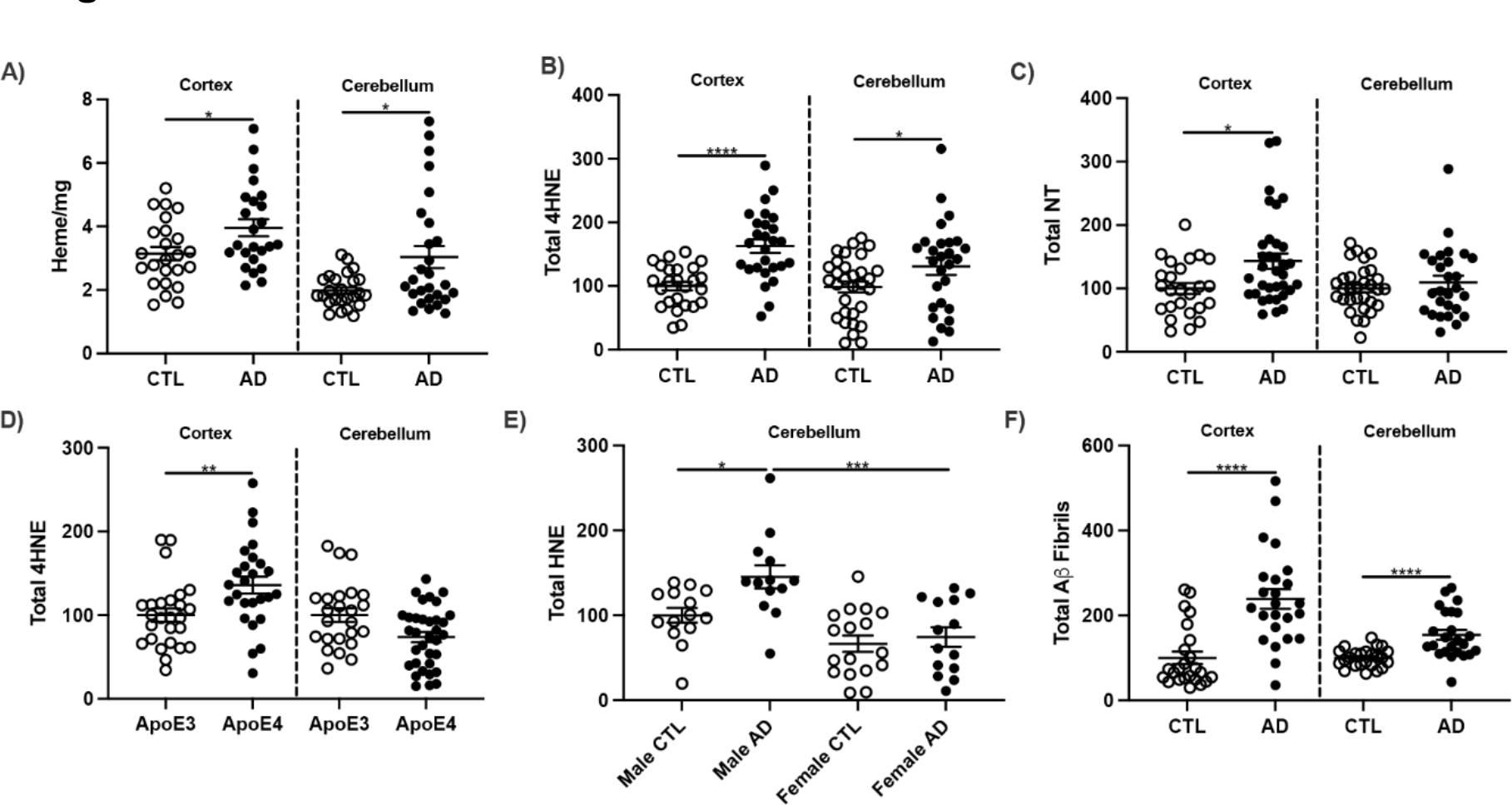
Oxidative damage (HNE, NT) in human prefrontal cortex and cerebellar proteins in age matched AD and cognitively normal control (CTL). Prefrontal cortex from Brodmann area 8, 9, or 10 and cerebellum were washed for measurements of **A)** heme, **B)** dot blot for 4HNE, **C)** dot blot for NT, **D)** HNE, **E)** HNE, **F)** total amyloid fibrils. Significance, 2-tailed t-test (A-D,F), one-way ANOVA with Tukey’s posthoc test (E): *p<0.05, **p<0.01, ***p<0.001, ****p<0.0001.

**Extended Data Figure 2:**
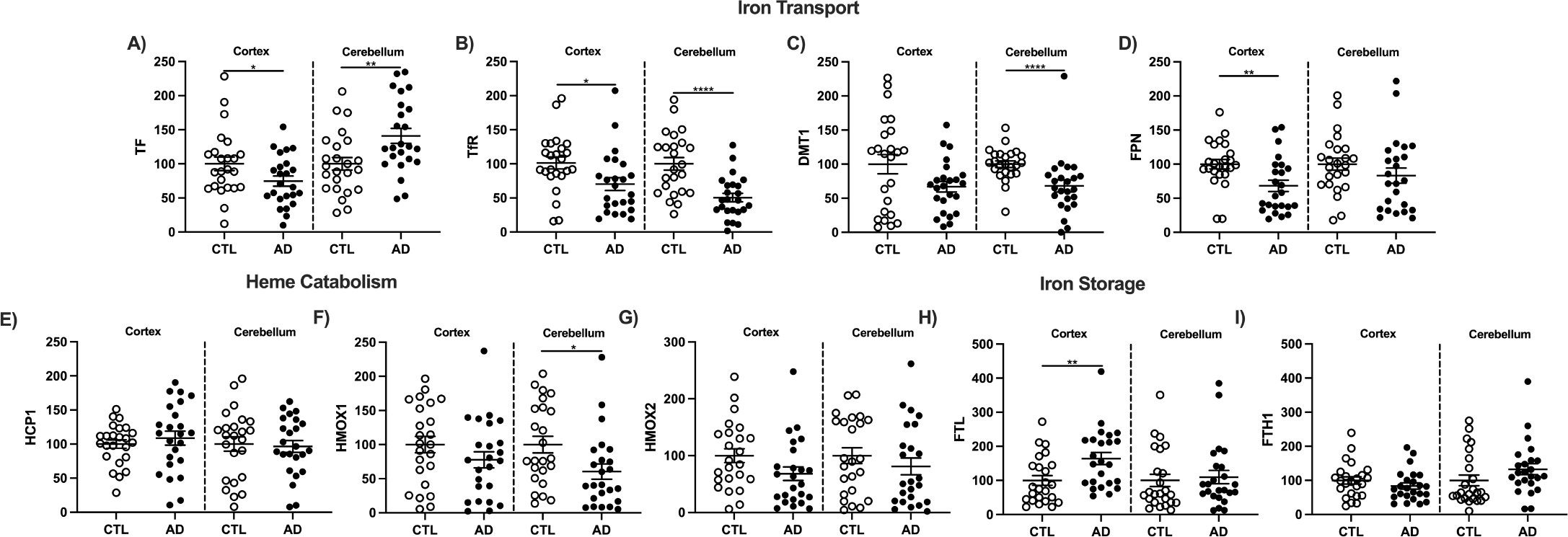
Iron transport and metabolism in AD vs cognitively normal prefrontal cortex and cerebellum. Western blots data for **A)** transferrin (TF), **B)** transferrin receptor (TfR/CD71), **C**) divalent metal transporter 1 (DMT1), **D)** ferroportin (FPN), **E)**, heme carrier protein 1 (HCP1), **F)** hemeoxygenase 1 (HMOX1), **G)** heme oxygenase 2 (HMOX2), **H)** ferritin light chain (FTL), **I)** ferritin heavy chain 1 (FTH1). Significance, 2-tailed t-test (A-I): *p<0.05, **p<0.01, ****p<0.0001.

**Extended Data Figure 3:**
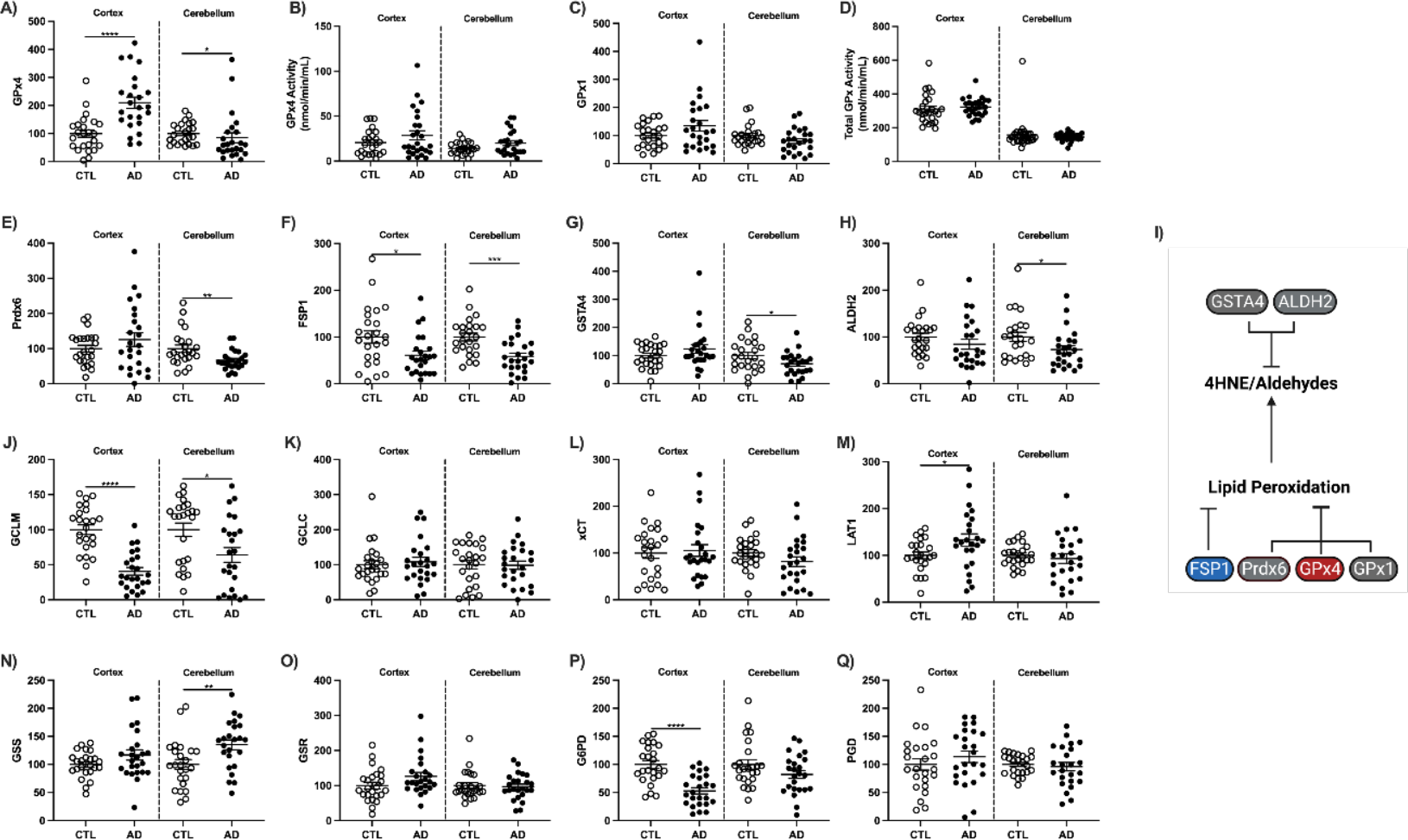
Enzymes that mitigate lipid peroxidation and facilitate GSH production. Western blot or enzyme activity units for prefrontal cortex or cerebellum from whole tissue lysate for **A)** GPx4, **B)** GPx4 activity, **C)** GPx1, **D)** total GPx activity, **E)** Prdx6, **F)** FSP1, **G)** GSTA4, and **H)** ALDH2. **I)** Enzymatic repair of lipid peroxidation. GSH cycle enzyme levels by Western blots, as relative fluorescent units (RFU) **J)** GCLM, **K)** GCLC, **L)** xCT/SLC7A11, **M)** LAT1, **N)** GSS, **O)** GSR, **P)** G6PD, and **Q)** PGD. Significance, 2-tailed t-test (A-H,J-Q): *p<0.05, **p<0.01, ***p<0.001, ****p<0.0001.

**Extended Data Figure 4:**
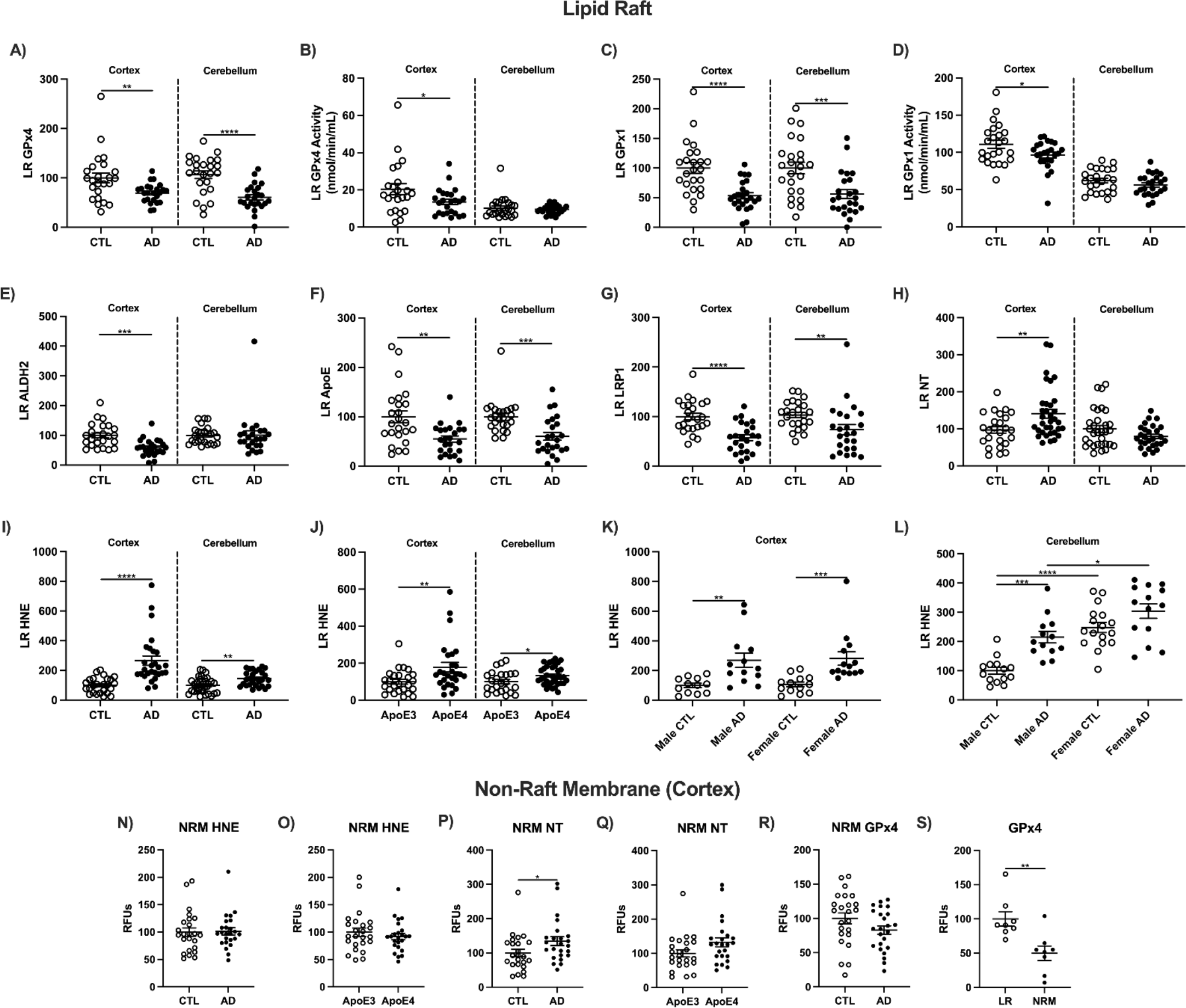
The AD lipid raft has increased damage and reduced antioxidant defense. **A)** Schema of lipid raft damage, protective mechanisms, and cholesterol shuttling in AD prefrontal cortex and cerebellum. Western blots as relative fluorescent units (RFUs) or enzymatic activity **B)** GPx4, **C)** GPx4 activity **D)** GPx1, **E)** GPx1 activity, **F)** ALDH2, **G)** ApoE, **H)** LRP1. Dot blots for **I)** NT, and HNE presented as **J)** CTL vs AD, **K)** ApoE allele, and sex for **L)** cortex and **M)** cerebellum. Significance, 2-tailed t-test (A-J, N-S), one-way ANOVA with Tukey’s posthoc test (K,L): *p<0.05, **p<0.01, ***p<0.001, ****p<0.0001.

**Extended Data Figure 5:**
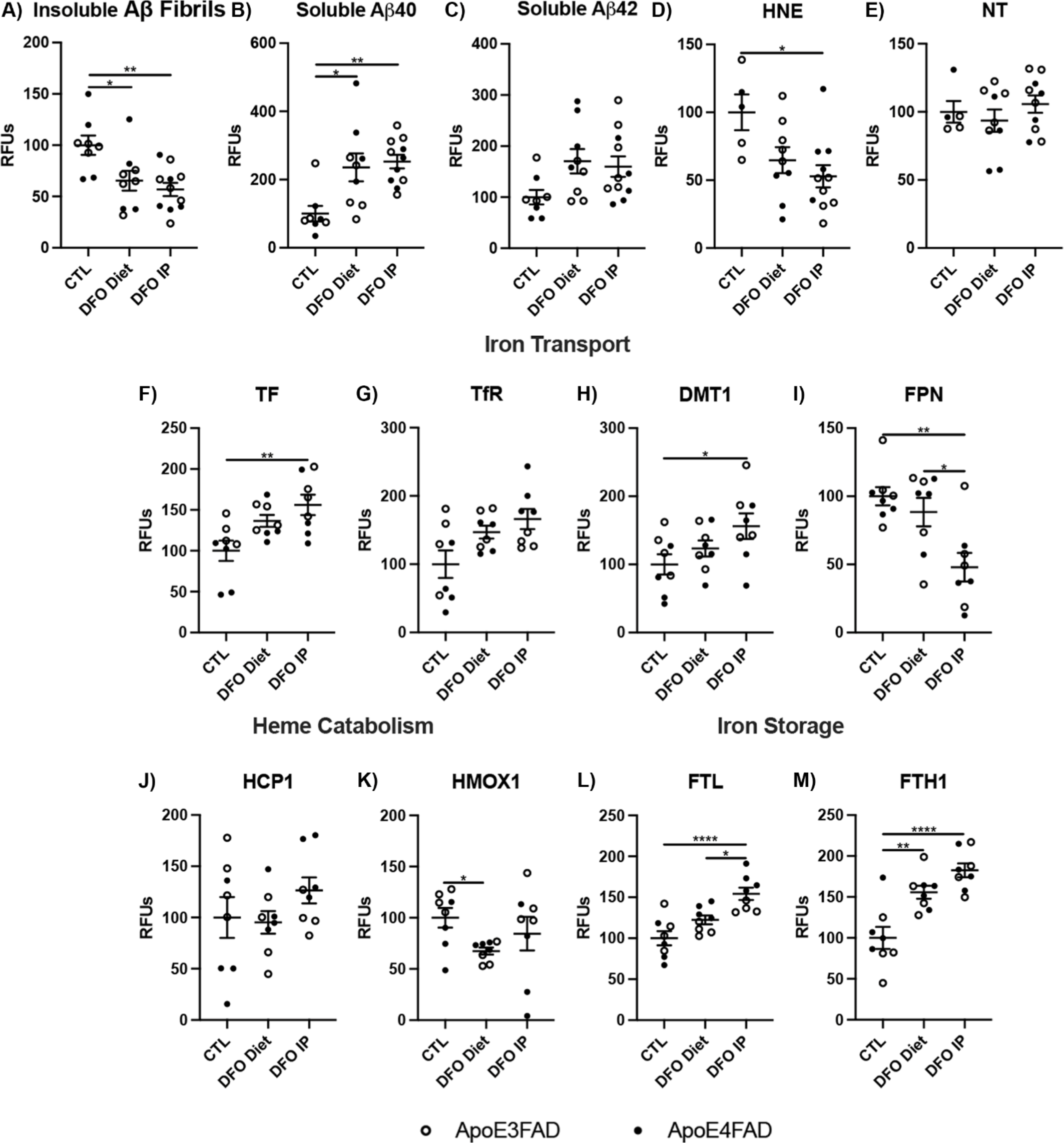
DFO chelation of EFAD mouse cortex comparing diet and IP: **A)** insoluble Aβ fibrils, **B)** soluble Aβ40, **C)** soluble Aβ42, **D)** HNE, **E)** NT, **F)** TF, **G)** TfR, **H)** DMT1, **I)** FPN, **J)** HCP1, **K)** HMOX1, **L)** FTL, and **M)** FTH1. Significance, 2-tailed t-test: *p<0.05, **p<0.01, ***p<0.001, ****p<0.0001.

**Extended Data Figure 6:**
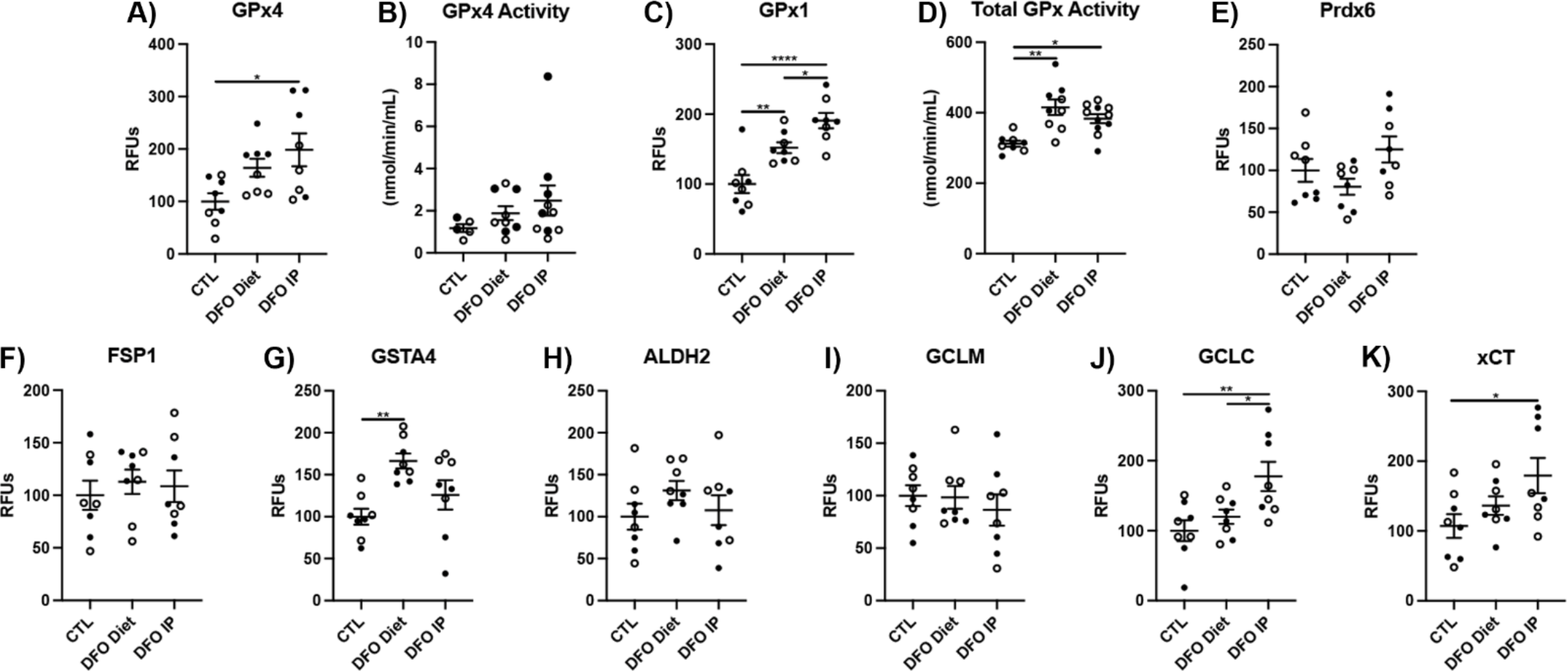
DFO chelation of EFAD mouse cortex comparing diet and IP: **A)** GPx4, **B)** GPx4 activity, **C)** GPx1, **D)** GPx1 activity, **E)** Prdx6, **F)** FSP1, **G)** GSTA4, **H)** ALDH2, **I)** GCLM, **J)** GCLC, **K)** xCT. Significance, one-way ANOVA with Tukey’s posthoc test (A-K): *p<0.05, **p<0.01, ***p<0.001, ****p<0.0001.

**Extended Figure 7:**
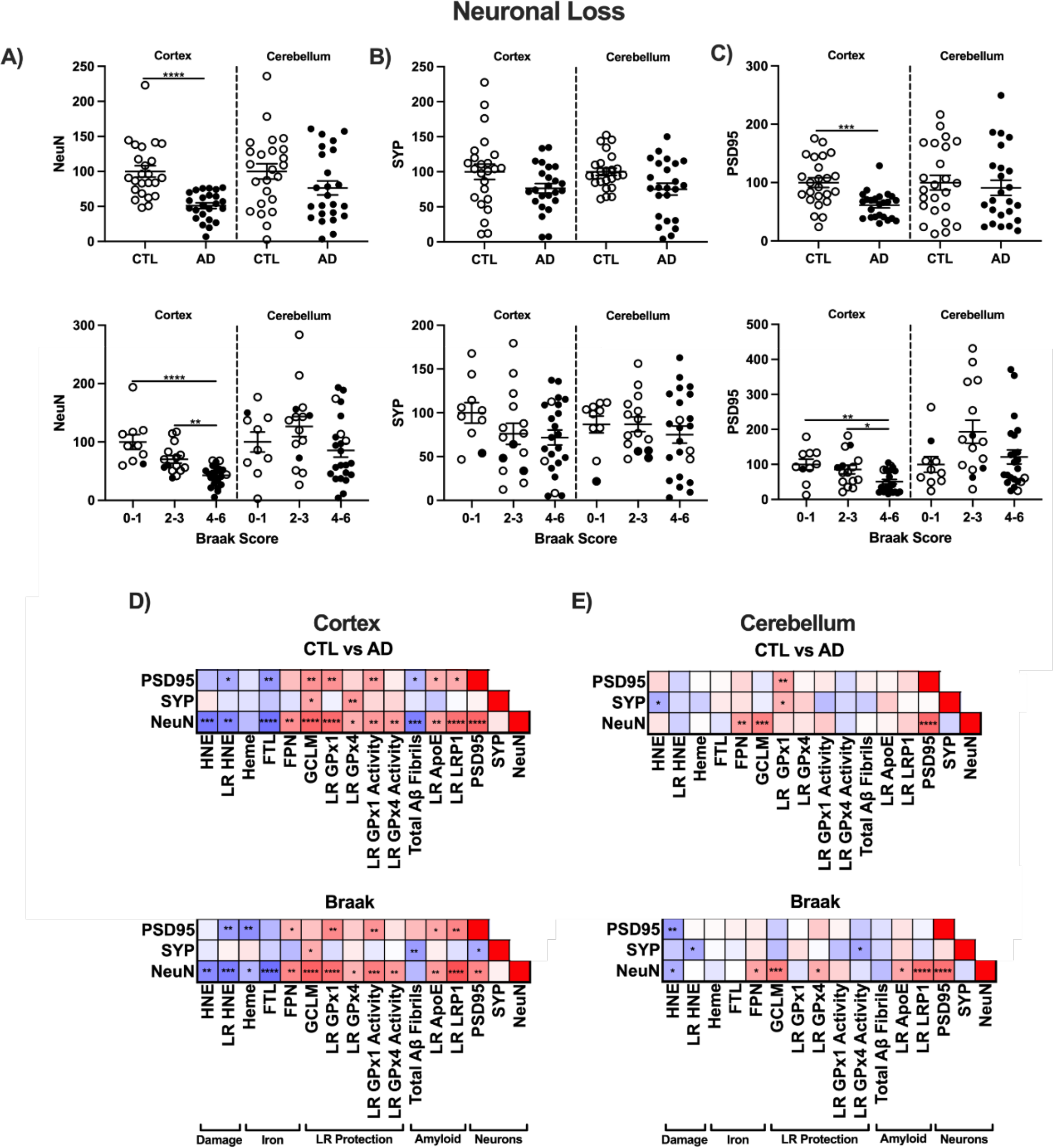
Neuronal loss in AD. Western blots as relative fluorescent units (RFUs) of neuronal markers shown by clinical CTL vs AD or Braak stage^41^. **A)** neuronal nuclei antigen (NeuN), **B)** synaptophysin 1 (SYP), **C)** postsynaptic density protein 95 (PSD95). Correlation matrixes for variables with significant relationships to NeuN for **D)** prefrontal cortex and **E)** cerebellum. Significance by 2- tailed t-test (CTL vs AD), One-way ANOVA (Braak) with Tukey’s posthoc, or by Spearman correlation: *p<0.05, **p<0.01, ***p<0.001, ****p<0.0001.

**Extended Figure 8:**
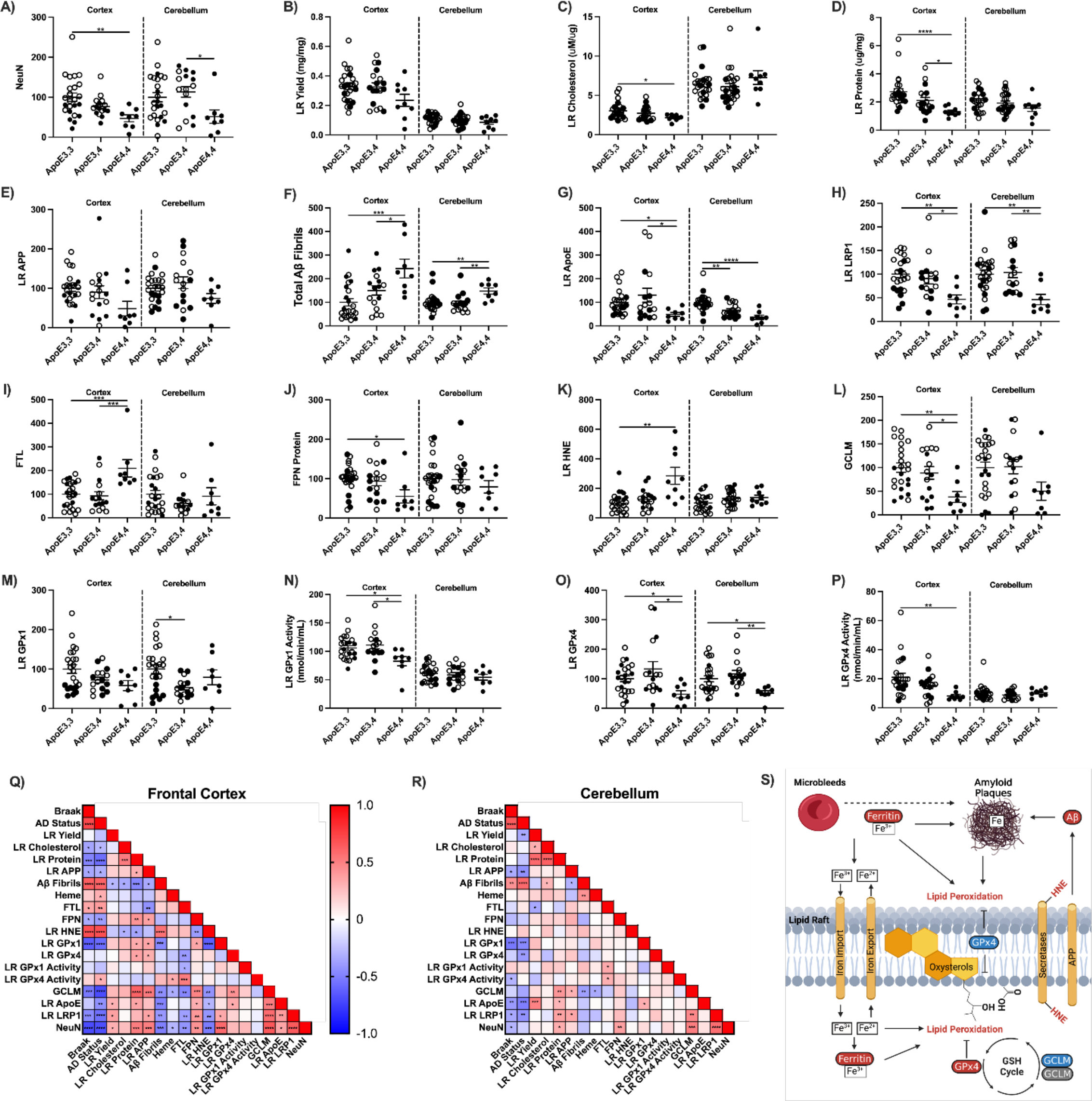
ApoE allele differences in lipid rafts, antioxidant defense, and amyloid. Levels of **A)** NeuN, **B)** LR yield, **C)** LR cholesterol, **D)** total LR protein, **E)** LR APP, **F)** total Aβ fibrils, **G)** LR ApoE, **H)** LR LRP1, **I)** FTL, **J)** FPN, **K)** LR HNE, **L)** GCLM, **M)** LR GPx1, **N)** LR GPx1 activity, **O)** LR GPx4, and **P)** LR GPx4 activity in prefrontal cortex and cerebellum of cognitively normal (open circles) and demented (closed circles). Correlation matrix of proteins analyzed by ApoE allele in **Q)** frontal cortex and **R)** cerebellum. **S)** Schematic hypothesis linking oxidized lipid rafts to amyloid processing. One way ANOVA with Tukey’s posthoc, or by Spearman correlation: *p<0.05, **p<0.01, ***p<0.001, ****p<0.0001.

